# Three Accessory Gene Clusters Drive Host-Adaptation in Group B *Streptococcus*

**DOI:** 10.1101/2023.08.10.552778

**Authors:** Chiara Crestani, Taya L. Forde, John Bell, Samantha J. Lycett, Laura M.A. Oliveira, Tatiana C.A. Pinto, Claudia G. Cobo-Ángel, Alejandro Ceballos-Márquez, Nguyen N. Phuoc, Wanna Sirimanapong, Swaine L. Chen, Dorota Jamrozy, Stephen D. Bentley, Michael Fontaine, Ruth N. Zadoks

## Abstract

*Streptococcus agalactiae* (Group B *Streptococcus*, GBS) is a major pathogen of humans and animals, posing a threat to human health as well as food security. Here, we investigate the role of genomic mechanisms, including homologous recombination and horizontal gene transfer, in shaping the population structure of GBS and its adaptation to three major host groups (humans, cattle, fishes). We demonstrate that the GBS population comprises host-specialist, host-adapted lineages as well as host generalists, and that these categories differ in their level or recombination. Although the accessory genome at large varies by lineage rather than host, genome wide association studies show that host association is driven by three accessory genome clusters, regardless of lineage or breadth of the host spectrum. These genomic clusters (*scpB* in human GBS, lactose operon in bovine GBS, Locus 3 in fish GBS) are known (*scpB*, Lac.2) or shown here (Locus 3) to be functionally relevant and are shared with other streptococcal species occupying the same host niche. These findings demonstrate the importance of considering the role of non-human host species in the evolution of GBS, including high risk clones that may lead to interspecies transmission and affect efficacy of future GBS vaccines.

## 1 Introduction

*Streptococcus agalactiae*, or Group B *Streptococcus* (GBS), is a major cause of neonatal infectious disease, with outcomes ranging from full recovery to neurodevelopmental impairment and mortality (Kohli-Lynch et al., 2017; Sinha et al., 2016). It also contributes to maternal disease, preterm birth, and stillbirth (Bianchi-Jassir et al., 2017; Hall et al., 2017; Seale et al., 2017). In 2015, the World Health Organisation identified “development of GBS vaccines suitable for maternal immunization in pregnancy and use in low- and middle-income countries” as a priority (WHO,2021). In non-pregnant adults, GBS can cause sepsis, pneumonia, urinary tract infection, skin and soft tissue infection, meningitis, and respiratory disease, with age, surgery, or comorbidities as predisposing factors (Collin et al., 2019; Navarro-Torné, Curcio, Moïsi, & Jodar, 2021). In Southeast Asia, invasive GBS disease due to foodborne infection has been described, often accompanied by osteoarthritis (Barkham et al., 2019; Luangraj et al., 2022). Like other important human pathogens, GBS is also a commensal (Chaguza et al., 2020; Richardson et al., 2018). The reported prevalence of colonisation with GBS outside pregnancy ranges from 9% (urethra) to 26% (rectum), with variation by detection method, age, and continent (van Kassel et al., 2021). Oropharyngeal carriage in humans has been linked to contact with neonates (Roloff, Stepanyan, & Valenzuela, 2018) as well as contact with animals (Cobo-Ángel et al., 2019; Nielsen & Emanuelson, 2013). Group B streptococci were first described by Rebecca Lancefield, who isolated them from “certified milk” and milk of a cow with mastitis (Lancefield, 1933). *Streptococcus agalactiae* continues to be an important cause of bovine mastitis around the world, with negative impact on milk quality and quantity, cow health, and farmers’ livelihoods (Krishnamoorthy et al., 2021). It was largely eliminated from northern Europe, but re-emerged in the 21^st^ century (Crestani et al., 2021; Jørgensen et al., 2016; Katholm, Bennedsgaard, Koskinen, & Rattenborg, 2012), possibly due to host-species jumping (Crestani et al., 2021). GBS is also a major pathogen of tilapia, the world’s third most farmed fish species (FAO,2022). It was first described in poikilothermic species (fishes, frogs) in the 1980s (Amborski, Snider 3rd, Thune, & Culley Jr, 1983). Its subsequent emergence as major fish pathogen and its global dissemination have been linked to the expansion and intensification of aquaculture around that time (Barkham et al., 2019; Kawasaki et al., 2018).

Multi-host bacterial pathogens can adapt to host species or tissue types using diverse mechanisms, from point mutations (Viana et al., 2015) to the acquisition, through horizontal gene transfer (HGT), of accessory genome content that provides a survival advantage in the context of a particular host, e.g., the ability to evade the host immune system or the acquisition of new metabolic pathways in response to the availability of particular nutrients (Lowder et al., 2009; Richardson et al., 2018). For example, a transposon carrying *scpB*, encoding a C5a-peptidase, which enables invasion of epithelial cells, has been associated with human GBS (Shabayek & Spellerberg, 2018; Sørensen, Poulsen, Ghezzo, Margarit, & Kilian, 2010). It has also been described in other human-pathogenic streptococci (e.g., *Streptococcus pyogenes* and *Streptococcus dysgalactiae* subsp. *equisimilis*), whereby interspecies transfer may be facilitated by its location on a mobile genetic element (MGE) (Davies et al., 2005; Franken et al., 2001; Gleich-Theurer et al., 2009). Nutritional adaptation is exemplified by the acquisition of an MGE encoding a lactose-fermenting pathway, which appears to be critical for successful colonisation of the bovine mammary gland (Richards et al., 2011). Considering the high intraspecific genome plasticity of GBS (Richards et al., 2019), and the emergence of new clades in new host species in the last hundred years (Barkham et al., 2019; Crestani et al., 2021; Da Cunha et al., 2014), there is a need to gain a better understanding of GBS host adaptation mechanisms, not least to ensure that emergence of new variants can be monitored during the anticipated introduction of human GBS vaccination (WHO,2021).

Here, we analyse the GBS population using 1,254 genomes, rigorously selected to represent diversity in lineages, host species and geographic origins, to gain insight into the role of genome plasticity and the accessory genome in GBS host adaptation. To facilitate this analysis, we propose nomenclature based on clonal groups (CG) and sublineages (SL), as recently introduced for other multi-host pathogens, and demonstrate that they comprise host generalists, host-adapted and host specialist lineages. Using genome-wide association studies (GWAS), we show that human, bovine and fish-adaptation of GBS are associated with C5a-peptidase (*scpB*), the lactose operon (Lac.2) and an accessory gene cluster known as Locus 3, respectively, regardless of GBS lineage, and provide functional evidence for the critical role of the latter in fish-associated GBS.

## 2 Results and Discussion

### 2.1 The global GBS population is composed of host-generalist and host-specialist lineages

The GBS global population comprises: i) host specialist lineages, whose occurrence is highly associated (*≥*98%) or restricted to a single host species (human, bovine, camel) or host group (poikilothermic species, including fishes and frogs; for simplicity, this host group is referred to as “fish” throughout the manuscript); ii) host-adapted lineages, which have a strong host predilection, defined here as having more than 80% but less than 98% of isolates originating from a single host or host group (see Supplementary appendix section A.3); iii) host generalists, defined here as having no more than 80% of isolates originating from a single host or host group (Fig. C.1).

Of 15 sub-populations that were identified by hierarchical Bayesian clustering, largely in alignment with the topology of the core gene phylogeny (Fig. 1A), six were host specialists, i.e. sublineage (SL) 22 (human), SL61, SL91 (bovine), SL552 (poikilothermic species), SL609 and SL612 (camel); four were host-adapted SL, i.e. SL17, SL19, SL26 and SL130 (all with a predilection for the human host); and the remaining five were host generalists, i.e. SL1, SL23, SL103, SL283 and SL314. In this nomenclature, sublineages are named after the most common sequence type (ST) in each subpopulation based on 7-gene multi-locus sequence typing (MLST) nomenclature (Fig. C.2). Within each SL, clonal groups (CG, n=23) were defined based on phylogenetic sub-clusters using the same nomenclature principle as for SL (Fig. C.2). Among CG, the same cut-offs used for SL identified eight host specialists, six host-adapted CGs, and nine host generalists (Fig. 1B and Fig. C.1). Two generalist SL (SL1, SL283) comprised three generalist CG each (CG1, CG459 and CG817; CG6, CG7 and CG283, respectively), as did one specialist with host-restricted CG (SL552, including CG260, CG261 and CG552). However, SL23, which was classified as host generalist, comprised host generalist CG23 and host-adapted CG24, which is primarily found in humans. The two CG are associated with different capsular types: the human-adapted CG24 primarily comprises isolates from capsular type Ia, whereas CG23 is mostly associated with capsular type III, except for a human-associated subclade of type Ia and II isolates (for more detail, please refer to our Microreact project at microreact.org). A more in-depth discussion of each SL (including CG, capsular types, host species, accessory genes and pilus islands) can be found in section 2.6 and in the supplementary appendix at section B.4.

**Fig. 1.**
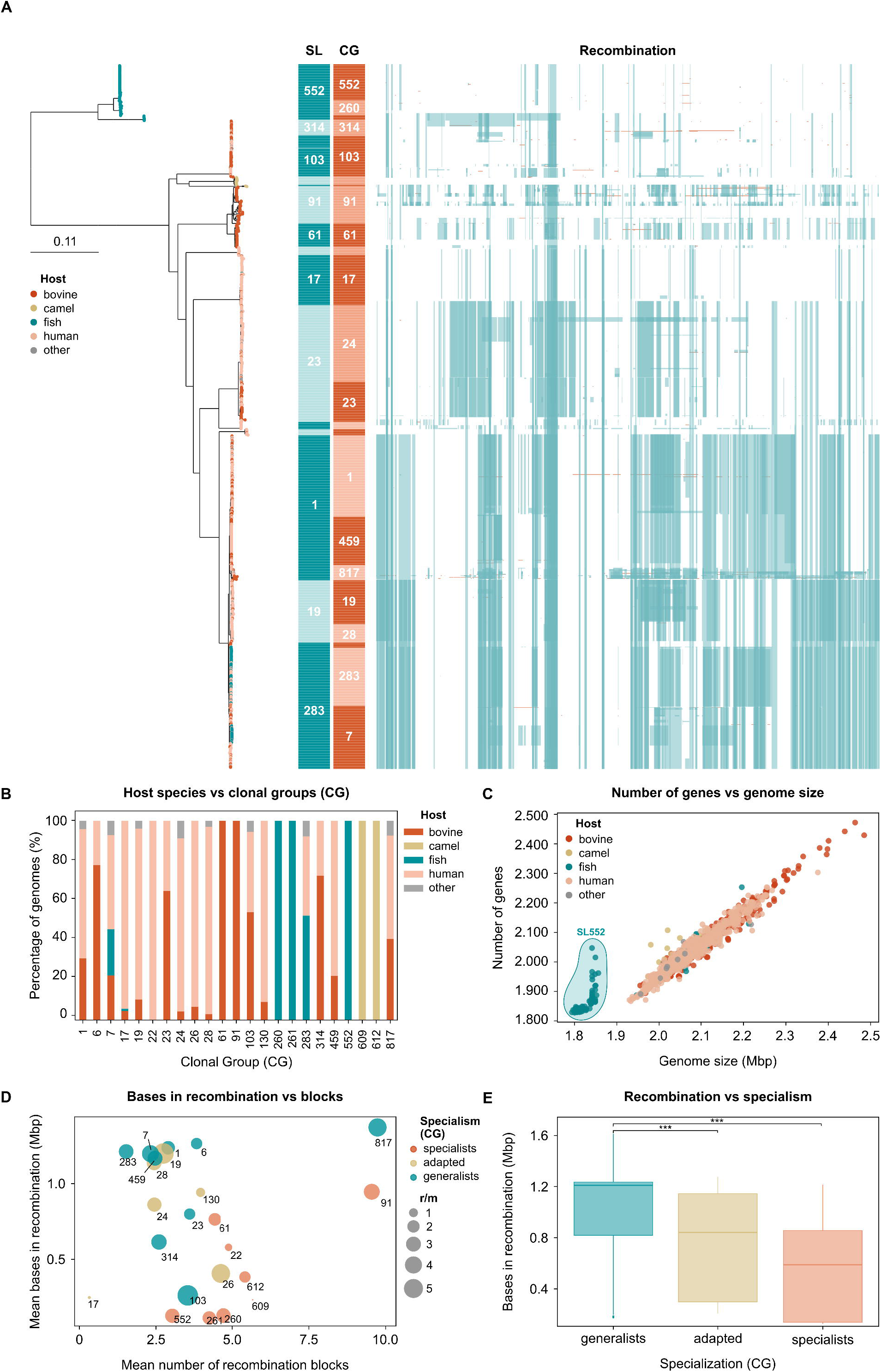
Population structure and pangenome of Group B *Streptococcus* (GBS). A) Maximum-likelihood phylogenetic tree of 1,254 GBS genomes; leaf colours indicate host of origin, and external strips show sublineage (SL), clonal group (CG) and homologous recombination; tree was rooted on an out-group of five reference genomes from *Streptococcus pyogenes* (hidden); B) Prevalence of host species within each CG; C) Correlation of number of genes and assembly size; SL552 shows a marked pseudogenisation and reduced genome size; D) Correlation of average number of recombination bases and recombination blocks of each CG, as well as recombination to mutation (r/m) rate; E) Recombination observed in the three categories of host-specialism; the difference between groups is statistically significant (p-value*<*0.0001).

The existence of host-generalist and host-specialist lineages within populations of multi-host bacterial pathogens is well-described, for example in *Salmonella enterica* (Langridge et al., 2015), *Campylobacter jejuni* (Sheppard et al., 2014, 2013), and *Staphylococcus aureus* (Lowder et al., 2009; Richardson et al., 2018). These observations have been associated with genomic phenomena that may lead to host-restriction, typically in the case of pseudogenisation and gene loss, as described for SL61 and SL552 (Almeida et al., 2016; Rosinski-Chupin et al., 2013), (see also section 2.5), or to host jumps, which provide access to a new accessory gene pool, with subsequent host-adaptation, which is often a result of acquisition of accessory genes that provide an adaptive advantage to the new ecological niche.

### 2.2 The accessory gene set in GBS isolates is not host-associated but lineage-associated

Network analysis of accessory gene distances shows distinct clusters that align with SL (and therefore with CG) rather than host species (Fig. 2), with the exception of host specialist lineages, such as SL552 or SL91, for which host species and lineage were here inherently fully concordant (100% prevalence of one host). These observations suggest that the accessory gene set as an ensemble depends more on the sublineage of origin of the isolates than on their ecological niche in GBS. This is in contrast to what is reported in *S. aureus* by Richardson et al. (2018), who showed an association between host species and the whole set of accessory genes. These results, coupled with previous knowledge on the association of mobile genetic elements encoding host-associated virulence factors or metabolic pathways (e.g., *scpB* in humans (Franken et al., 2001; Gleich-Theurer et al., 2009) and Lac.2 in bovines (Lyhs et al., 2016; Richards, Choi, Bitar, Gurjar, & Stanhope, 2013; Richards et al., 2011)), led us to hypothesise that a limited number of acquired genes might drive host-adaptation.

**Fig. 2.**
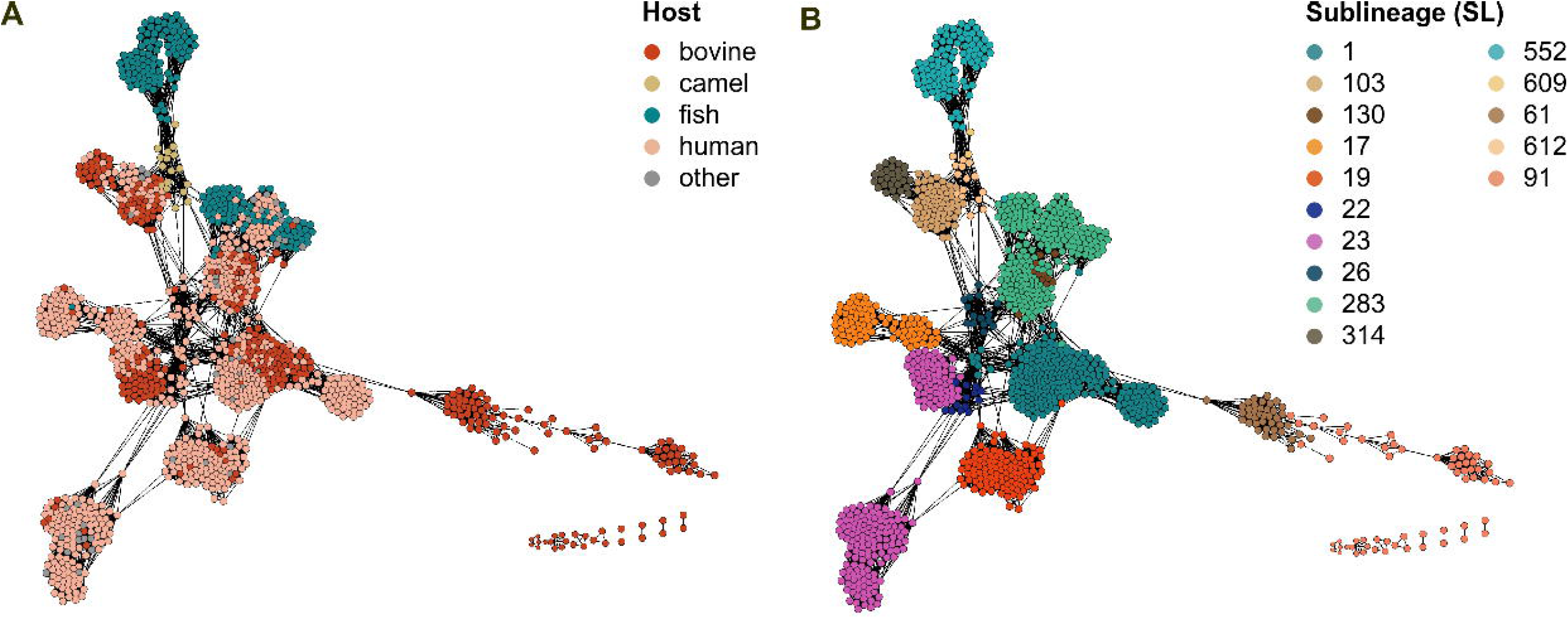
Accessory gene distance network of 1,254 Group B *Streptococcus* (GBS) genomes. A) Major host groups (human, bovine, fish and camel) are overlaid to the nodes; B) Sublineages (SL) defined by fastbaps and renamed based on an inheritance principle from 7-gene MLST nomenclature are shown. The two panels show association of accessory gene clusters with SL, and a lack of clustering based on host species, unless when this is a direct result of host-specific lineages (e.g., SL552, SL91, SL61).

### 2.3 Three accessory gene clusters are highly host-associated in GBS independently of lineage

We performed GWAS with two approaches, one pangenome-based and the other unitig-based, to detect association of accessory genes with the three major GBS host groups. Both analyses show that three accessory gene clusters, each associated with one host group, dominate all other genes for their significant positive association.

#### GBS in humans: the *scpB* transposon

In human GBS, the pangenome-based approach identified the *scpB* transposon as significantly positively associated (*p*-value for *scpB* was 1.26×10^−131^; all *p*-values reported in this paper for pan-GWAS analyses were corrected with the Benjamini-Hochberg method), and similar results were obtained with the unitig-based approach (Fig. 3A, Fig. C.3A). The *scpB* gene is a transposon-encoded virulence factor which is known for its higher prevalence among human isolates (Franken et al., 2001; Sørensen et al., 2010), although its association with this host group had never been described before as a result of GWAS in such a vast and representative GBS dataset. This gene interacts negatively with the human host immune system (cleaving the C5a complement component), it contributes to GBS cellular adhesion and invasion, and it has been associated with the ability to colonise or infect the human host in other human pathogenic streptococci (Group A *Streptococcus*, *Streptococcus dysgalactiae* subsp. *equisimilis* and *Streptococcus canis*) (Franken et al., 2001; Gleich-Theurer et al., 2009). *In vitro* work suggests that *scpB* likely plays no role in bovine GBS infections (Gleich-Theurer et al., 2009), even when it is carried by GBS cattle isolates. Its role in fish infections has not been assessed. Interestingly, in our dataset, *scpB* in GBS from fish was uniquely found in a lineage shared with humans (SL283), which could explain why this is the only lineage from fish that shows evidence for zoonotic transmission (see section 2.6.1).

**Fig. 3.**
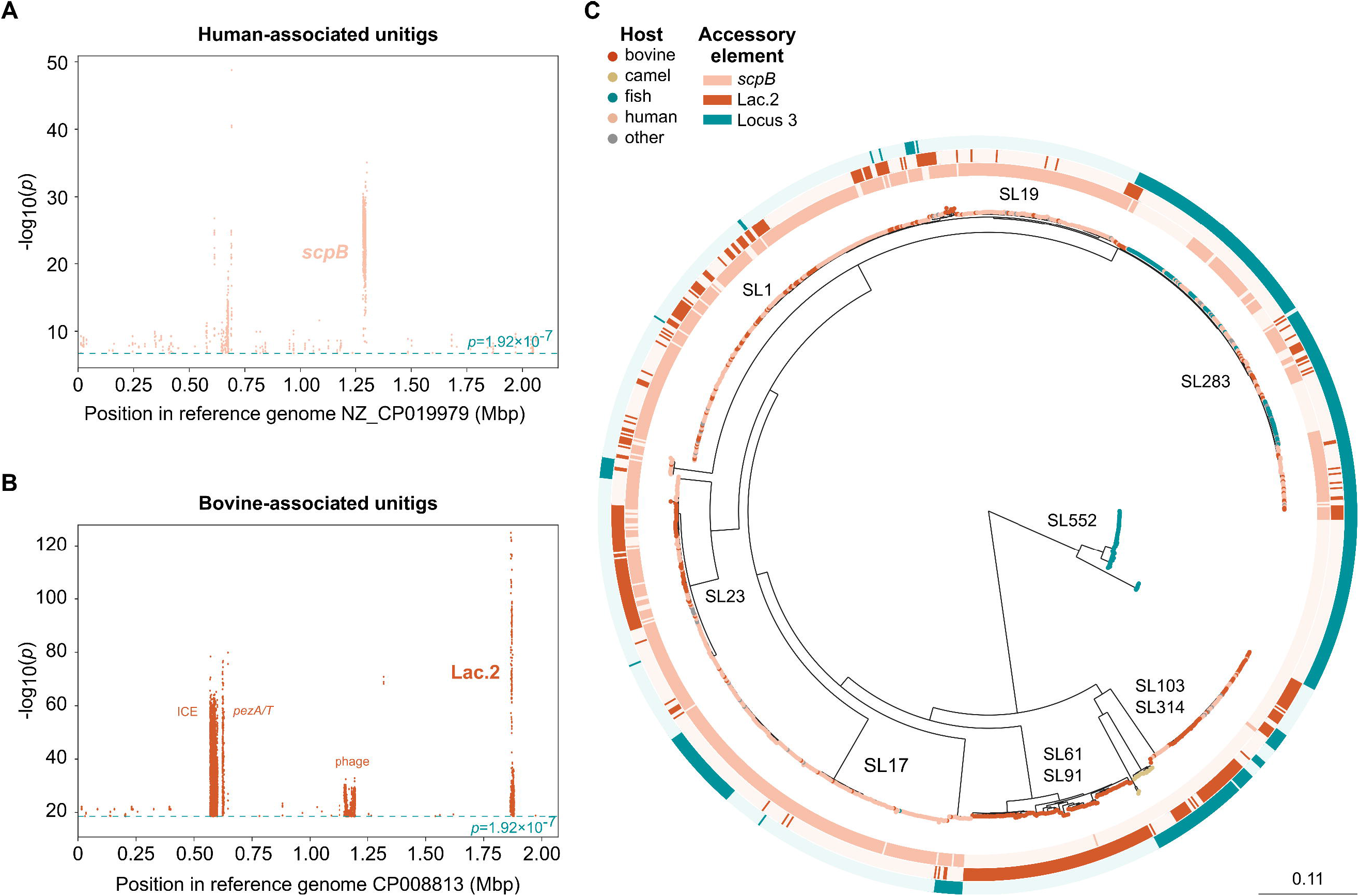
Host-associated accessory gene clusters in Group B *Streptococcus* (GBS). A) Manhattan plot of human-associated unitigs mapped to reference genome NZ CP019979; B) Manhattan plot of bovine-associated unitigs mapped to reference genome CP008813; C) Circular maximum-likelihood phylogenetic tree of 1,254 GBS genomes. Leaf colours indicate host of origin, whereas the three external strips show presence/absence of the three main host-associated accessory gene clusters (human-*scpB*, bovine-Lac.2, fish-Locus 3, respectively).

#### GBS in animals: the Lac.2 operon and Locus 3

In bovine GBS, significant genes with both GWAS approaches corresponded to a 9 to 11-gene cluster, the Lac.2 operon, in particular its genes *lacEG* (Scoary *p*-value 4.69×10^−168^; Fig. 3B, Fig. C.3B). A Lac.2+ genetic background is associated with phenotypic lactose fermentation, which has been observed primarily in cattle GBS isolates (Lyhs et al., 2016; Richards et al., 2013). In cattle, GBS only causes infections localised to the mammary gland, which is rich in lactose. The acquisition of metabolic pathways (here: Lac.2) in response to nutrient availability (here: lactose) is a known driver of niche adaptation (Holt et al., 2015; Sheppard et al., 2013; Sheppard, Guttman, & Fitzgerald, 2018). The detection of Lac.2 in other bovine mastitis-causing streptococci (e.g., *Streptococcus dysgalactiae* subsp. *dysgalactiae* and *Streptococcus uberis* (Richards et al., 2013, 2011)) strengthens the argument that acquisition of lactose-fermenting genes drives niche adaptation to the bovine mammary gland.

In fish GBS, the unitig-based approach was unsatisfactory, which was likely due to a strong population structure effect, as genomes from this host group are found only in two SL (SL552 and SL283). With the pangenome-based approach, genes belonging to a cluster known as Locus 3 scored the highest based on sensitivity, specificity and *p*-values (SE 99.5%; SP 72.6%; BH *p*-value 9.62×10^−85^). Locus 3, which is described as fish-associated (Delannoy et al., 2016), was present in 99.5% of fish and frog GBS genomes and only in a minority of non-fish assemblies, especially in the generalist SL283, a SL primarily shared between humans and fish. The only exception was an ST17 isolate attributed to fish. Unlike for *scpB* in humans and Lac.2 in bovines, there is no evidence for a functional role of Locus 3 in fish (see section 2.4). However, we detected a segment of this element in all available genomes (NCBI, February 2023; data not shown) from another important agent of fish streptococcosis, *Streptococcus iniae* (Agnew & Barnes, 2007), highlighting its association with the aquatic niche and its possible impact in streptococcal adaptation to fish. The significant association of the three accessory gene clusters with the three main GBS host groups across SL and CG (Fig. 3C) as well as with other streptococci pathogenic to the same host species, suggests that they are each major drivers of adaptation to the distinct host niches in GBS lineages, and possibly more broadly in the genus *Streptococcus*.

### 2.4 Locus 3 is essential for infection in fish in various GBS genetic backgrounds

Locus 3 is a 17-gene cluster inserted between a GNAT family N-acetyltransferase and a class I SAM-dependent methyltransferase, which includes, among others, genes for carbohydrate transport and metabolism (Delannoy et al., 2016) (Fig. 4A). Nile tilapia (*Oreochromis niloticus*) challenge experiments were carried out to compare virulence of a multi-gene knock-out mutant (ΔLocus3) with wildtype (WT) isolates in two distinct, highly virulent fish-associated GBS backgrounds (ST7 and ST283). Animals were either challenged intraperitoneally (IP) or through cohabitation with the IP-challenged fish. For both challenge routes and both GBS strains, a significant reduction in mortality was observed in groups challenged with the ΔLocus3 mutant relative to WT (Tab. 1, Fig. 4B and Fig. 4C). The observed attenuation of GBS after removal of Locus 3 provides the first functional evidence of its importance in GBS infection of fish.

**Fig. 4.**
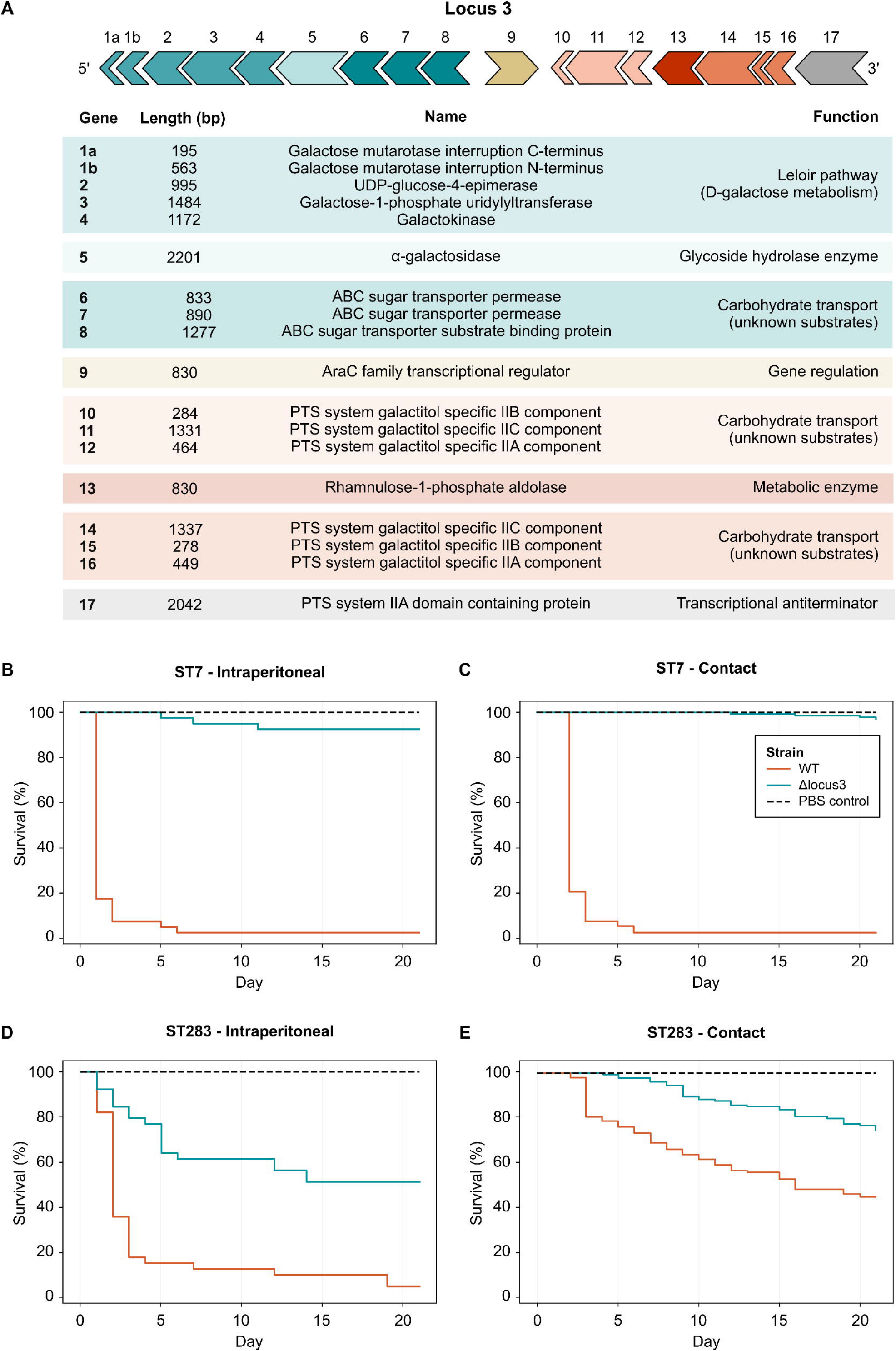
Locus 3 gene cluster and survival curves of Nile tilapia (*Oreochromis niloticus*). A) Locus 3 gene cluster organisation and associated gene functions; B) Tilapia challenged intraperitoneally (IP) with Group B *Streptococcus* (GBS) Sequence Type (ST) 7 wild type (WT) or its isogenic mutant (ΔLocus3); C) Tilapia challenged with GBS ST7 (WT) or its isogenic mutant (ΔLocus3) through cohabitation with IP-challenged fish; D) Tilapia challenged IP with GBS ST283 (WT) or its isogenic mutant (ΔLocus3); E) Tilapia challenged with GBS ST283 (WT) or its isogenic mutant (ΔLocus3) through cohabitation with IP-challenged fish. All experiments (B-E) included a negative control (mock challenge with phosphate buffered saline).

**Table 1.** Tilapia mortality analysis. Relative mortality of Nile tilapia (*Oreochromis niloticus*) challenged with Group B *Streptococcus* (GBS) sequence type (ST) 7 and ST283 mutants (ΔLocus3), compared to the wildtype (WT) strain. Fish were challenged intraperitoneall (IP) or through cohabitation with IP-challenged fish.

| Strain | IP challenged |  |  | Cohabitation challenged |  |  |
| --- | --- | --- | --- | --- | --- | --- |
|  | EMP<sup>1</sup> | RM<sup>2</sup> | SA<sup>3</sup> | EMP | RM | SA |
| ST7 WT | 95 |  |  | 88 |  |  |
| ST7 ΔLocus3 | 0 | -100 | 5.4x10<sup>-7**</sup> | 4 | -96 | 3.4x10<sup>-22***</sup> |
| ST283 WT | 95 |  |  | 56 |  |  |
| ST283 ΔLocus3 | 25 | -74 | 5.3x10<sup>-6**</sup> | 16 | -71 | 5.3x10<sup>-8**</sup> |

<sup>1</sup>End point mortality (EPM) expressed in percentage (%). <sup>2</sup>Relative mortality (RM) expressed in percentage (%). <sup>3</sup>Survival analysis (SA) *p*-value.

### 2.5 Host-specialism among GBS lineages is associated with different levels of genome plasticity

In addition to the acquisition of accessory genome content, other genomic phenomena may affect levels of host-specialisation (generalists, adapted, specialists). For example, gene loss and pseudogenisation cause host-restriction in several bacterial pathogens, including *Salmonella enterica* subsp. *Enterica* serovar Gallinarum and Pullorum in poultry (Langridge et al., 2015), and for *Staphylococcus aureus* CC133 in ruminants (Guinane et al., 2010). In GBS, this mechanism is observed in the host-restricted SL552, which has marked pseudogenisation (Richards et al., 2019) and a reduced genome size (Fig. 1C). Recombination can also play a role in host-adaptation (Sheppard et al., 2018). We observed marked differences in recombination between clonal groups of GBS, ranging from absence in e.g., CG552, CG260, CG261, to large recombination blocks in e.g., CG283, CG19, CG817 (Fig. 1A and Fig. 1D). Host-generalist lineages appear more subject to recombination than host-specialists, as illustrated by the number of nucleotide bases present within recombination blocks, for which the difference is statistically significant (t-value generalists=107.56, adapted=-15.42, specialists=-28.12, p-value*<*0.0001) (Fig. 1E).

Our results are indicative of a higher genome plasticity of host-generalist lineages compared to host-specialists and suggest that the ability to uptake and retain foreign DNA, including accessory genes that could provide a survival advantage in new niches and hosts, is superior in host-generalists (e.g., SL283). The lack of shared recombination between some lineages (e.g., SL1, SL19 and SL283 vs others; SL23 vs others; SL91 vs others, SL61 vs others), coupled with results from analysis of the accessory gene set (section 2.2), suggests the existence of some barriers to genetic exchange (i.e., HGT) within GBS. These barriers are unlikely to be ecological (e.g., segregation (Mourkas et al., 2022)), as at least some lineages co-exist in the different host populations and in the same geographical areas (Cobo-Ángel et al., 2019; Crestani et al., 2021). Rather, these are more likely mechanistic barriers that can act as a defense against the uptake of foreign DNA, such as restriction-modification systems (RMS), CRISPR, or antiphage systems, as described by Mourkas et al. (2022).

### 2.6 The impact of host-associated accessory gene carriage on public health: examples in CG283 and SL23

#### 2.6.1 CG283

CG283 (Fig. 5A) is widespread in fishes in the South-East Asian region (Barkham et al., 2019), carried asymptomatically by close to 3% of the population in some areas (Barkham et al., 2023), and it is the causative agent of the first ever reported outbreak of GBS foodborne disease. All CG283 isolates in our study carried the fish-associated Locus 3. With regards to *scpB*, traits analysis predicted it to be present in the common ancestor of CG283 isolates and to have been lost multiple times, with two major events corresponding to two branches (the first around 1992, and the second around 2002) (Fig. 5A). The presence of *scpB* in the ancestral node of CG283 suggests a probable human origin for this clade, which would have acquired Locus 3 concomitantly to the expansion of aquaculture (1980s). We hypothesise that the exchange of accessory genetic material between human and fish GBS is favoured by frequent direct or indirect contact between humans and fish in South-East Asia, e.g., through recycling of human waste as fish feed, floating fish farms or fish handling and consumption. A CG283 sub-clade mostly carrying *scpB* (Fig. 5A) is associated with human infections, and it includes genomes from GBS isolated during the 2015 Singaporean foodborne outbreak that was linked with the consumption of contaminated raw fish (Barkham et al., 2019; Kalimuddin et al., 2017). In this sub-clade, *scpB* appears to have been lost independently at least six times. One of these loss events, which happened around 2002 (Fig. 5A), appears to have taken place in Vietnam, and was followed by expansion of a CG283 sub-clade primarily associated with fish. The emergence of CG283 in Brazil, estimated to have occurred in 2015 and already described in 2016 (Leal, Queiroz, Pereira, Tavares, & Figueiredo, 2019), likely resulted from introduction of the Vietnamese sub-clade (Fig. 5A). So far, no human infections associated with CG283 have been reported in South America. This is probably not a result of differences in eating habits compared to the Asian population, as the introduction of Japanese-style cuisine and raw fish consumption in Brazil dates back to the beginning of the 1900s. Rather, absence of reported CG283 infections in humans could be due to the absence of *scpB*. The evolution of CG283 illustrates how ongoing surveillance of GBS across host species is needed to detect emergence of new clades as well as the acquisition and loss of accessory genome content that informs on the hazard to public health and food security.

**Fig. 5.**
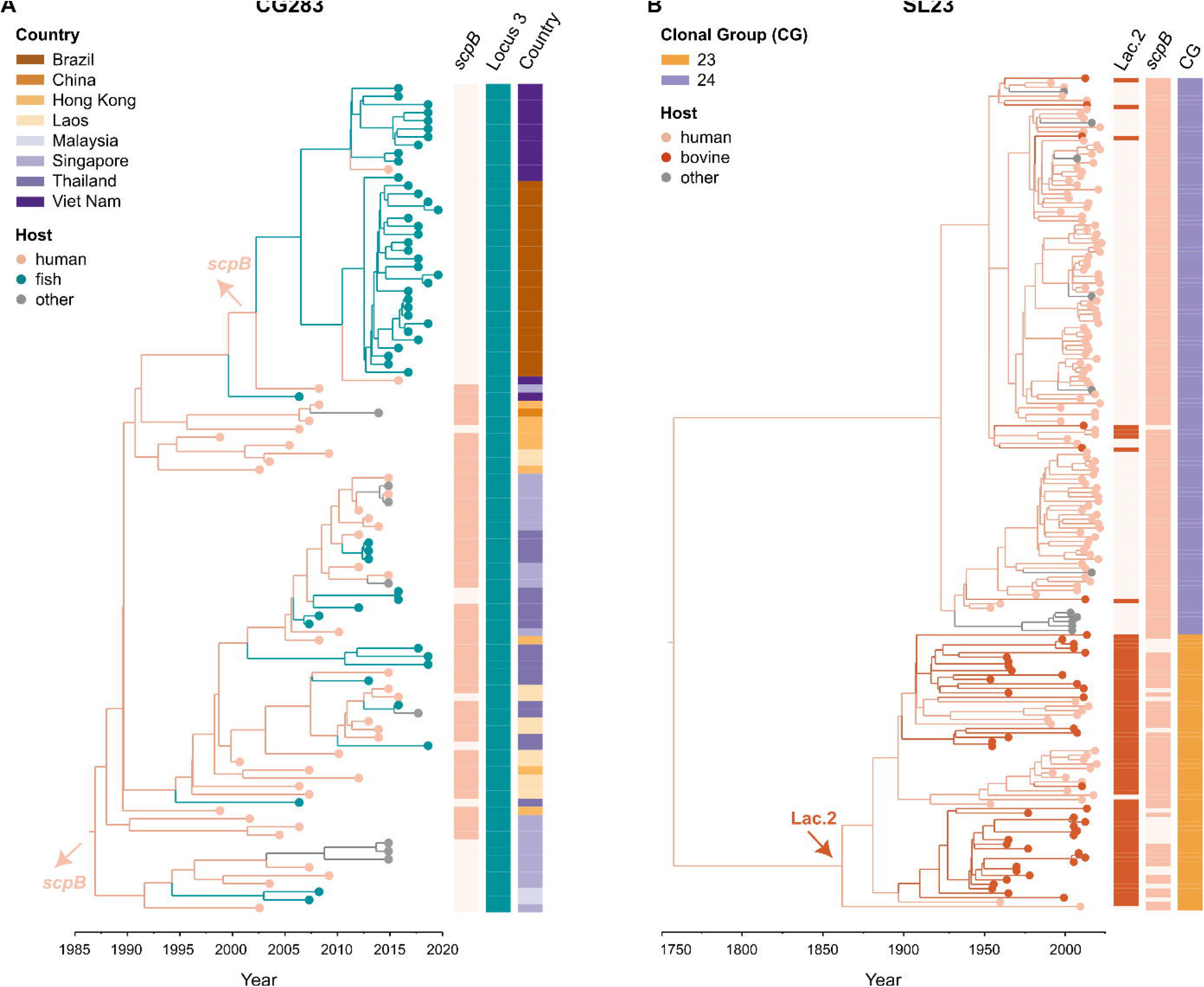
Time-scaled phylogenies of two clones of public health interest in Group B *Streptococcus* (GBS). Branch colour shows host-jumps, leaf colour the host of isolation, and external strips show presence-absence of genes and features of interest. A) Clonal group (CG) 283 is predicted to have originated around 1986 from humans. Its population shows diversification in multiple sub-clades over the years. Genomes associated with human cases from the 2015 outbreak of foodborne infection in Singapore are part of a sub-clade in which most of the genomes carry the human-associated transposon bearing *scpB*. In recent years, CG283 has been described in Brazil (first isolates available are from 2015); this introduction likely happened from Vietnam, and it is associated with a *scpB* - genotype. The two main *scpB* loss events are indicated with arrows on the phylogeny; B) Sublineage (SL) 23 likely originated from humans at the beginning of the 18^th^ century. It is described as a host-generalist clonal complex (CC). However, when assessing host prevalence within its two CG (CG23 and CG24), as well as the distribution of human and bovine-associated accessory gene clusters (*scpB* and Lac.2, respectively) the two CG show distinct patterns of host-tropism. CG24 appears as human-adapted, whereas isolates of CG23, whose common ancestor is predicted to have acquired Lac.2 in the second half of the 1800s (red arrow), show a tendency towards bovine-specialism, but retain the ability to cause human infections (*scpB* is highly conserved).

#### 2.6.2 SL23

SL23, which was referred to as CC23 in previous literature, is an old GBS sublineage, with emergence estimated to have occurred in the middle of the 18^th^ century in our study and by others (Da Cunha et al., 2014). It has long been considered a host-generalist but there are two CG within SL23, with distinct patterns in terms of host tropism. The first one, CG24, is a host-adapted CG that primarily causes disease in humans (Fig. 1), with occasional host jumps to cattle. These host jumps are associated with sporadic acquisitions of Lac.2 (Fig. 5B). Reflecting its human host association, *scpB* is strongly conserved in this CG, with the exception of one bovine isolate. The second CG in SL23 is CG23, which is a host-generalist clade that primarily affects humans and cattle, with additional reports in, e.g., companion animals, aquatic mammals and reptiles (Delannoy et al., 2013). Mirroring the situation for *scpB* in CG24, Lac.2 is highly conserved in CG23, with the exception of one human isolate (Fig. 5B), explaining its ability to infected the bovine mammary gland. As in CG283, *scpB* has been lost multiple times independently from CG23, suggesting that this virulence factor is lost easily in the absence of selective pressure from the human host. Considering that the phylogeny suggests bidirectional interspecies transmission of CG23 (Fig. 5B) ongoing monitoring of CG23 infections in both host species would be needed to gauge risk of bovine GBS to public health.

## 3 Conclusions

Multi-host bacterial pathogens often comprise lineages that show distinct patterns of host-tropism, which can vary from host generalism, such as CC130 and CC398 in *S. aureus* (Richardson et al., 2018), to host restriction, such as for the serovars Gallinarum and Pullorum of *S. enterica* subsp. *enterica* in poultry (Langridge et al., 2015). A multitude of inter-playing ecological (e.g., physical segregation), genomic (e.g., ability of a genome to uptake DNA, homologous and non-homologous recombinations, gene loss and pseudogenisation) and adaptive mechanisms can affect host-range (Mourkas et al., 2022). Population gene introgression and genome plasticity within GBS have been described as major driving forces in the evolution of GBS (Richards et al., 2019). Our results show that several genomic mechanisms, including recombination and pseudogenisation, have shaped the GBS population and the host association of its sublineages, with higher genome plasticity in host-generalists than in host specialists.

As in other bacterial pathogens (Mourkas et al., 2022; Richardson et al., 2018), adaptive mechanisms that confer a fitness advantage within the context of a particular niche play a crucial role in GBS. Despite the diversity of lineages associated with each major host category, and the diversity of host categories associated with many major lineages, we have identified three key accessory gene clusters that are major driving forces of GBS host-adaptation, namely *scpB* in human, Lac.2 in bovine, and Locus 3 in fish. Of note, we describe functional evidence for the critical role of Locus 3 in fish infections in different genetic backgrounds of GBS. Furthermore, our study builds upon existing evidence for the role of *scpB* in human infections (Gleich-Theurer et al., 2009) and Lac.2 in bovine infections (Richards et al., 2011), further solidifying the significance of these genomic elements in host adaptation. Remarkably, these three elements are not only present in GBS but are also found in other streptococcal species that share ecological niches with GBS (e.g., *scpB* in Group A *Streptococcus*, *Streptococcus dysgalactiae* subsp. *equisimilis* and *Streptococcus canis*, Lac.2 in *S. dysgalactiae* subsp. *dysgalactiae* and *S. uberis*, and Locus 3 in *S. iniae*), suggesting their broader importance within the genus *Streptococcus*.

Through analysis of two multihost clades (CG283 and SL23), we show the impact of acquisition or loss of these three elements on host-adaptation and expansion following host-jumps, and illustrate the public health relevance of such genomic mechanisms for GBS. We propose that, by focusing genomic surveillance on clones with high potential for host-jumps (i.e., those with high levels of recombination, such as SL283 and SL1) and their acquisition of hostassociated accessory gene clusters, strategies and interventions to reduce the risk of interspecies transmission could be improved.

## 4 Materials and Methods

A detailed description of all materials and methods can be found in the Supplementary appendix file, section A.

### 4.1 Genome selection

First, we aimed to collate a dataset representative of the broad diversity of the GBS population in terms of clonal complexes, hosts of origin, geographical locations, and temporal range (n=2,437). Metadata were curated from the literature (March 2020) and whole genome sequence data were obtained either from public repositories or self-generated. Using Pandas v1.1.3 (The Pandas Development Team, 2020), we de-duplicated the initial dataset for clones that were over-represented based based on a series of metadata variables (see section A.2) and further filtered based on genome assembly quality (see Supplementary appendix). The resulting de-duplicated, quality-filtered dataset comprised 1,254 genomes.

### 4.2 Core genome analyses

IQ-TREE v.2.0.6 (Nguyen, Schmidt, Von Haeseler, & Minh, 2015) was used to estimate a core genome phylogeny with a GTR model, from a recombination-free alignment file obtained with snippy v4.4.5 (https://github.com/tseemann/snippy) and gubbins v3.2.0 (Croucher et al., 2014), using NGBS128 as reference. On the same alignment file, fastbaps v1.0.8 (Tonkin-Hill, Lees, Bentley, Frost, & Corander, 2019) was used to define genomic clusters.

A generalised linear model was run in RStudio v2022.07.01, R v4.2.0 (2022-04-22), on the output from gubbins, after having mapped the internal nodes and the leaves to the corresponding CG, to test association between the host-specialisation level (generalists, adapted, specialists) and the number of nucleotide bases identified in recombination blocks.

### 4.3 Accessory genome analyses

Accessory gene distances were calculated with GraPPLE (https://github.com/JDHarlingLee/GraPPLE) using the Jaccard similarity index from a gene presence/absence matrix; the latter was generated with panaroo v1.2.0 (Tonkin-Hill et al., 2020) from gff files annotated with Prokka v1.14.5 (Seemann, 2014). The resulting file was visualised with Graphia v2.2 (Freeman et al., 2022).

### 4.4 Genome-Wide Association Studies

Scoary v1.6.16 (Brynildsrud, Bohlin, Scheffer, & Eldholm, 2016) was run to detect gene-enrichment in GBS from the three major host groups from the panaroo-generated presence/absence gene matrix. The host groups (human, bovine, and the poikilotherm group including fishes and frogs) were defined as a binary phenotypes, with 1 as belonging to the host, and 0 as not belonging to the host; GBS genomes from host groups other than these three were always categorised as 0, including those originating from dead fish sampled at markets for which a possible contamination due to human handling could not be excluded.

The pyseer suite v1.3.3 (Lees, Galardini, Bentley, Weiser, & Corander, 2018) was used to assess unitig association with a linear mixed model on the same host groups as above.

### 4.5 *In vivo* assessment of the role of Locus 3 in fish infection

Challenge studies on Nile tilapia (*Oreochromis niloticus*) were carried out with ST7 and ST283 knock-out mutants of Locus 3 (ΔLocus3). Strains and plasmids used for mutagenesis are described in the Supplementary appendix (Tab. C.1). In preparation for challenge, tilapia were transferred into 100 L experimental tanks (4 replicates per GBS strain, 2 negative control replicates) under the same ecological conditions. For challenge experiments, ten fish from each experimental tank were challenged by intraperitoneal (IP) injection with 0.1 mL of phosphate buffered saline (PBS; negative control) or 0.1 mL of 10^5^ CFU/mL of GBS in PBS. For contact challenge experiments, IP-challenged tilapia were co-habited with 40 healthy tilapia per tank.

Mortality of IP-challenged and contact-challenged fish was monitored for 21 days. Moribund fish were euthanized and included in mortality counts. Kidney and brain samples were taken from up to 10 fish per tank and used to check for presence of GBS (bacterial culture) and Locus 3 genes (PCR assays).

Kaplan-Meier survival analysis (May, 2009) was conducted to compare survival of knock-out mutants and their isogenic wild type strains. A statistical Log-rank test was conducted (Kleinbaum & Klein, 2012), using Microsoft Excel software, to compare survival curves of tilapia challenged by wild type vs knock-out mutants.

### 4.6 Time-scaled phylogenies of two clones of public health interest: CG283 and SL23

BEAST v2.6.6 (Bouckaert et al., 2014) was used to estimate the time of emergence of two clones of public health interest, CG283 and SL23. The best model parameters chosen were: GTR+G4 with a strict clock and a constant coalescent population size. Xml files were run in triplicate with 200 million generations and sampling frequency of 10,000 (burn-in 10%). Resulting trees from the three replicate runs were combined with LogCombiner, and the final tree files were obtained with TreeAnnotator and visualised with FigTree v1.4.4 (http://tree.bio.ed.ac.uk/software/figtree/) and Microreact (Argimón et al., 2016).

The Maximum Clade Credibility trees from the two BEAST datasets were used together with gene presence/absence for *scpB*, Lac.2, and Locus 3 in a discrete traits analyses, as well as with host. To infer the gene presence/absences at the ancestral nodes in the trees, the ancestral character estimation function (ace) within R-package ape (Paradis & Schliep, 2019) was used with a discrete asymmetric model (all rates different, ARD).

## Supplementary information

Supplementary material, including detailed methodologies, results, figures and tables can be found in the Supplementary appendix file, sections A, B and C.

Supplementary table S1 gathers together genome metadata (both duplicated and de-duplicated datasets), assembly quality metrics, results from genomic typing and bioinformatic analyses. Supplementary tables S2, S3, and S4 show results from Scoary analyses for the human, bovine and fish phenotypes, respectively.

## Acknowledgments

We would like to thank John Lees for bioinformatic support for pyseer. We would also like to thank Marta Mansos-Lourenco, Carla Parada-Rodrigues and Sylvain Brisse for constructive feedback on the manuscript.

## Declarations

### Availability of data and materials

All raw or assembled Illumina sequence data is available from the European Nucleotide Archive (ENA) or from the National Center for Biotechnology Information (NCBI) (accession numbers can be found in the supplementary metadata file). Phylogenetic analysis of the 1,254 Group B *Streptococcus* genomes, together with all metadata, is available at the project URL https://microreact.org/project/vkEcchaHsSJa6sLhSGvmHF-group-b-streptococcus-host-adaptation-2023 within Microreact.

### Authors’ contributions

CC and RZ conceived and designed the study; LO, TP, CCA, AC, WS, NNP, SLC, DJ and SB collected bacterial isolates, epidemiological data, and sequence data. JB performed mutagenesis under supervision of MF. SL performed traits analysis and CC conducted all other data analyses and visualisation. CC, TF and RZ drafted the manuscript. All authors read and approved the manuscript.

## References

Agnew, W., & Barnes, A.C. (2007). *Streptococcus iniae*: an aquatic pathogen of global veterinary significance and a challenging candidate for reliable vaccination. Veterinary Microbiology, 122 (1-2), 1–15.

Almeida, A., Alves-Barroco, C., Sauvage, E., Bexiga, R., Albuquerque, P., Tavares, F., … Glaser, P. (2016). Persistence of a dominant bovine lineage of group B *Streptococcus* reveals genomic signatures of host adaptation. Environmental Microbiology, 18 (11), 4216–4229.

Amborski, R., Snider 3rd, T., Thune, R., Culley Jr, D. (1983). A non-hemolytic, group b *Streptococcus* infection of cultured bullfrogs, rana catesbeiana, in brazil. Journal of Wildlife Diseases, 19 (3), 180–184.

Argimón, S., Abudahab, K., Goater, R.J., Fedosejev, A., Bhai, J., Glasner, C., … others (2016). Microreact: visualizing and sharing data for genomic epidemiology and phylogeography. Microbial Genomics, 2 (11), e000093.

Barkham, T., Tang, W.Y., Wang, Y.-C., Sithithaworn, P., Kopolrat, K.Y., Worasith, C. (2023). Human fecal carriage of *Streptococcus agalactiae* sequence type 283, thailand. Emerging Infectious Diseases, 29 (8), 1627–1629.

Barkham, T., Zadoks, R.N., Azmai, M.N.A., Baker, S., Bich, V.T.N., Chalker, V., … Chen, S.L. (2019). One hypervirulent clone, sequence type 283, accounts for a large proportion of invasive *Streptococcus agalactiae* isolated from humans and diseased tilapia in Southeast Asia. PLoS Neglected Tropical Diseases, 13 (6), e0007421.

Bianchi-Jassir, F., Seale, A.C., Kohli-Lynch, M., Lawn, J.E., Baker, C.J., Bartlett, L., … others (2017). Preterm birth associated with group b *Streptococcus* maternal colonization worldwide: systematic review and meta-analyses. Clinical Infectious Diseases, 65 (suppl 2), S133–S142.

Bouckaert, R., Heled, J., Kühnert, D., Vaughan, T., Wu, C.-H., Xie, D., … Drummond, A.J. (2014). Beast 2: a software platform for bayesian evolutionary analysis. PLoS Computational Biology, 10 (4), e1003537.

Brynildsrud, O., Bohlin, J., Scheffer, L., Eldholm, V. (2016). Rapid scoring of genes in microbial pan-genome-wide association studies with Scoary. Genome Biology, 17 (1), 1–9.

Chaguza, C., Senghore, M., Bojang, E., Gladstone, R.A., Lo, S.W., Tientcheu, P.-E., … others (2020). Within-host microevolution of *Streptococcus pneumoniae* is rapid and adaptive during natural colonisation. Nature communications, 11 (1), 3442.

Cobo-Ángel, C.G., Jaramillo-Jaramillo, A.S., Palacio-Aguilera, M., Jurado-Vargas, L., Calvo-Villegas, E.A., Ospina-Loaiza, D.A., … Ceballos-Marquez, A. (2019). Potential group B *Streptococcus* interspecies transmission between cattle and people in Colombian dairy farms. Scientific Reports, 9 (1), 1–9.

Collin, S.M., Shetty, N., Guy, R., Nyaga, V.N., Bull, A., Richards, M.J., … others (2019). Group b *Streptococcus* in surgical site and non-invasive bacterial infections worldwide: A systematic review and meta-analysis. International Journal of Infectious Diseases, 83, 116–129.

Crestani, C., Forde, T.L., Lycett, S.J., Holmes, M.A., Fasth, C., Persson-Waller, K., Zadoks, R.N. (2021). The fall and rise of group B *Streptococcus* in dairy cattle: reintroduction due to human-to-cattle host jumps? Microbial Genomics, 7 (9), 000648.

Croucher, N.J., Page, A.J., Connor, T.R., Delaney, A.J., Keane, J.A., Bentley, S.D., … Harris, S.R. (2014). Rapid phylogenetic analysis of large samples of recombinant bacterial whole genome sequences using gubbins. Nucleic acids research, 43 (3), e15–e15.

Da Cunha, V., Davies, M.R., Douarre, P.-E., Rosinski-Chupin, I., Margarit, I., Spinali, S., … Ma, L. (2014). *Streptococcus agalactiae* clones infecting humans were selected and fixed through the extensive use of tetracycline. Nature Communications, 5, ncomms5544.

Davies, M.R., Tran, T.N., McMillan, D.J., Gardiner, D.L., Currie, B.J., Sriprakash, K.S. (2005). Inter-species genetic movement may blur the epidemiology of streptococcal diseases in endemic regions. Microbes and Infection, 7 (9-10), 1128–1138.

Delannoy, C.M., Crumlish, M., Fontaine, M.C., Pollock, J., Foster, G., Dagleish, M.P., … Zadoks, R.N. (2013). Human *Streptococcus agalactiae* strains in aquatic mammals and fish. BMC Microbiology, 13 (1), 1–9.

Delannoy, C.M., Zadoks, R.N., Crumlish, M., Rodgers, D., Lainson, F.A., Ferguson, H., … Fontaine, M.C. (2016). Genomic comparison of virulent and non-virulent *Streptococcus agalactiae* in fish. Journal of Fish Diseases, 39 (1), 13–29.

Franken, C., Haase, G., Brandt, C., Weber-Heynemann, J., Martin, S., Lämmler, C., … Spellerberg, B. (2001). Horizontal gene transfer and host specificity of beta-haemolytic streptococci: the role of a putative composite transposon containing *scpB* and *lmb*. Molecular Microbiology, 41 (4), 925–935.

Freeman, T.C., Horsewell, S., Patir, A., Harling-Lee, J., Regan, T., Shih, B.B., … Angus, T. (2022). Graphia: A platform for the graph-based visualisation and analysis of high dimensional data. PLoS Computational Biology, 18 (7), e1010310.

Gleich-Theurer, U., Aymanns, S., Haas, G., Mauerer, S., Vogt, J., Spellerberg, B. (2009). Human serum induces streptococcal c5a peptidase expression. Infection and Immunity, 77 (9), 3817–3825.

Guinane, C.M., Ben Zakour, N.L., Tormo-Mas, M.A., Weinert, L.A., Lowder, B.V., Cartwright, R.A., … Fitzgerald, J.R. (2010). Evolutionary genomics of *Staphylococcus aureus* reveals insights into the origin and molecular basis of ruminant host adaptation. Genome Biology and Evolution, 2, 454–466.

Hall, J., Adams, N.H., Bartlett, L., Seale, A.C., Lamagni, T., Bianchi-Jassir, F., … others (2017). Maternal disease with group b *Streptococcus* and serotype distribution worldwide: systematic review and meta-analyses. Clinical infectious diseases, 65 (suppl 2), S112–S124.

Holt, K.E., Wertheim, H., Zadoks, R.N., Baker, S., Whitehouse, C.A., Dance, D., … Thomson, N.R. (2015). Genomic analysis of diversity, population structure, virulence, and antimicrobial resistance in *Klebsiella pneumoniae*, an urgent threat to public health. Proceedings of the National Academy of Sciences, 112 (27), E3574–E3581.

Jørgensen, H., Nordstoga, A., Sviland, S., Zadoks, R., Sølverød, L., Kvitle, B., Mørk, T. (2016). *Streptococcus agalactiae* in the environment of bovine dairy herds - rewriting the textbooks? Veterinary Microbiology, 184, 64–72.

Kalimuddin, S., Chen, S.L., Lim, C.T., Koh, T.H., Tan, T.Y., Kam, M., … Tang, W.Y. (2017). 2015 epidemic of severe *Streptococcus agalactiae* sequence type 283 infections in Singapore associated with the consumption of raw freshwater fish: a detailed analysis of clinical, epidemiological, and bacterial sequencing data. Clinical Infectious Diseases, 64 (suppl 2), S145–S152.

Katholm, J., Bennedsgaard, T., Koskinen, M., Rattenborg, E. (2012). Quality of bulk tank milk samples from Danish dairy herds based on real-time polymerase chain reaction identification of mastitis pathogens. Journal of Dairy Science, 95 (10), 5702–5708.

Kawasaki, M., Delamare-Deboutteville, J., Bowater, R.O., Walker, M.J., Beatson, S., Zakour, N.L.B., Barnes, A.C. (2018). Microevolution of *Streptococcus agalactiae* ST-261 from Australia indicates dissemination via imported tilapia and ongoing adaptation to marine hosts or environment. Applied and Environmental Microbiology, 84 (16), e00859–18.

Kleinbaum, D.G., & Klein, M. (2012). *Survival Analysis: A Self-Learning Text*. Springer, New York.

Kohli-Lynch, M., Russell, N.J., Seale, A.C., Dangor, Z., Tann, C.J., Baker, C.J., … others (2017). Neurodevelopmental impairment in children after group b streptococcal disease worldwide: systematic review and meta-analyses. Clinical Infectious Diseases, 65 (suppl 2), S190–S199.

Krishnamoorthy, P., Suresh, K.P., Jayamma, K.S., Shome, B.R., Patil, S.S., Amachawadi, R.G. (2021). An understanding of the global status of major bacterial pathogens of milk concerning bovine mastitis: A systematic review and meta-analysis (scientometrics). Pathogens, 10 (5), 545.

Lancefield, R.C. (1933). A serological differentiation of human and other groups of hemolytic streptococci. Journal of Experimental Medicine, 57 (4), 571–595.

Langridge, G.C., Fookes, M., Connor, T.R., Feltwell, T., Feasey, N., Parsons, B.N., … Thomson, N.R. (2015). Patterns of genome evolution that have accompanied host adaptation in *Salmonella*. Proceedings of the National Academy of Sciences, 112 (3), 863–868.

Leal, C.A., Queiroz, G.A., Pereira, F.L., Tavares, G.C., Figueiredo, H.C. (2019). *Streptococcus agalactiae* Sequence Type 283 in farmed fish, Brazil. Emerging Infectious Diseases, 25 (4), 776.

Lees, J.A., Galardini, M., Bentley, S.D., Weiser, J.N., Corander, J. (2018). Pyseer: a comprehensive tool for microbial pangenome-wide association studies. Bioinformatics, 34 (24), 4310–4312.

Lowder, B.V., Guinane, C.M., Zakour, N.L.B., Weinert, L.A., Conway-Morris, A., Cartwright, R.A., … Fitzgerald, J.R. (2009). Recent human-to-poultry host jump, adaptation, and pandemic spread of *Staphylococcus aureus*. Proceedings of the National Academy of Sciences, 106 (46), 19545–19550.

Luangraj, M., Hiestand, J., Rasphone, O., Chen, S.L., Davong, V., Barkham, T., … Keoluangkhot, V. (2022). Invasive *Streptococcus agalactiae* st283 infection after fish consumption in two sisters, lao pdr. Wellcome Open Research, 7, 148.

Lyhs, U., Kulkas, L., Katholm, J., Waller, K.P., Saha, K., Tomusk, R.J., Zadoks, R.N. (2016). *Streptococcus agalactiae* serotype IV in humans and cattle, northern Europe. Emerging Infectious Diseases, 22 (12), 2097.

May, W.L. (2009). Kaplan-Meier Survival Analysis. Encyclopedia of Cancer (p. 1590–1593). Springer, New York.

Mourkas, E., Yahara, K., Bayliss, S.C., Calland, J.K., Johansson, H., Mageiros, L., … others (2022). Host ecology regulates interspecies recombination in bacteria of the genus *Campylobacter*. eLife, 11, e73552.

Navarro-Torné, A., Curcio, D., Moïsi, J.C., Jodar, L. (2021). Burden of invasive group b *Streptococcus* disease in non-pregnant adults: A systematic review and meta-analysis. PloS one, 16 (9), e0258030.

Nguyen, L.T., Schmidt, H.A., Von Haeseler, A., Minh, B.Q. (2015). IQ-TREE: a fast and effective stochastic algorithm for estimating maximum-likelihood phylogenies. Molecular Biology and Evolution, 32 (1), 268–274.

Nielsen, C., & Emanuelson, U. (2013). Mastitis control in Swedish dairy herds. Journal of Dairy Science, 96 (11), 6883–6893.

Paradis, E., & Schliep, K. (2019). ape 5.0: an environment for modern phylogenetics and evolutionary analyses in r. Bioinformatics, 35 (3), 526–528.

Richards, V.P., Choi, S.C., Bitar, P.D.P., Gurjar, A.A., Stanhope, M.J. (2013). Transcriptomic and genomic evidence for *Streptococcus agalactiae* adaptation to the bovine environment. BMC Genomics, 14 (1), 920.

Richards, V.P., Lang, P., Bitar, P.D.P., Lefébure, T., Schukken, Y.H., Zadoks, R.N., Stanhope, M.J. (2011). Comparative genomics and the role of lateral gene transfer in the evolution of bovine adapted *Streptococcus agalactiae*. Infection, Genetics and Evolution, 11 (6), 1263–1275.

Richards, V.P., Velsko, I.M., Alam, T., Zadoks, R.N., Manning, S.D., Pavinski Bitar, P.D., … Stanhope, M.J. (2019). Population gene introgression and high genome plasticity for the zoonotic pathogen *Streptococcus agalactiae*. Molecular Biology and Evolution, 36 (11), 2572–2590.

Richardson, E.J., Bacigalupe, R., Harrison, E.M., Weinert, L.A., Lycett, S., Vrieling, M., … Fitzgerald, J.R. (2018). Gene exchange drives the ecological success of a multi-host bacterial pathogen. Nature Ecology and Evolution, 2 (9), 1468.

Roloff, K., Stepanyan, G., Valenzuela, G. (2018). Prevalence of oropharyngeal group b *Streptococcus* colonization in mothers, family, and health care providers. PloS one, 13 (9), e0204617.

Rosinski-Chupin, I., Sauvage, E., Mairey, B., Mangenot, S., Ma, L., Da Cunha, V., … Glaser, P. (2013). Reductive evolution in *Streptococcus agalactiae* and the emergence of a host adapted lineage. BMC Genomics, 14 (1), 252.

Seale, A.C., Bianchi-Jassir, F., Russell, N.J., Kohli-Lynch, M., Tann, C.J., Hall, J., … Bartlett, L. (2017). Estimates of the burden of group B streptococcal disease worldwide for pregnant women, stillbirths, and children. Clinical Infectious Diseases, 65, S200–S219.

Seemann, T. (2014). Prokka: rapid prokaryotic genome annotation. Bioinformatics, 30 (14), 2068–2069.

Shabayek, S., & Spellerberg, B. (2018). Group b streptococcal colonization, molecular characteristics, and epidemiology. Frontiers in microbiology, 9, 437.

Sheppard, S.K., Cheng, L., Méric, G., De Haan, C.P., Llarena, A.-K., Marttinen, P., … Jukka, C. (2014). Cryptic ecology among host generalist *Campylobacter jejuni* in domestic animals. Molecular Ecology, 23 (10), 2442–2451.

Sheppard, S.K., Didelot, X., Meric, G., Torralbo, A., Jolley, K.A., Kelly, D.J., … Falush, D. (2013). Genome-wide association study identifies vitamin B5 biosynthesis as a host specificity factor in *Campylobacter*. Proceedings of the National Academy of Sciences, 110 (29), 11923–11927.

Sheppard, S.K., Guttman, D.S., Fitzgerald, J.R. (2018). Population genomics of bacterial host adaptation. Nature Reviews Genetics, 19 (9), 549–565.

Sinha, A., Russell, L., Tomczyk, S., Verani, J., Schrag, S., Berkley, J., … Kim, S. (2016). Gbs vaccine cost-effectiveness analysis in sub-saharan africa working group. disease burden of group b *Streptococcus* among infants in sub-saharan africa: a systematic literature review and meta-analysis. Pediatr Infect Dis J, 35 (9), 933–42.

Sørensen, U.B.S., Poulsen, K., Ghezzo, C., Margarit, I., Kilian, M. (2010). Emergence and global dissemination of host-specific *Streptococcus agalactiae* clones. MBio, 1 (3), e00178–10.

The Pandas Development Team (2020, February). pandas-dev/pandas: Pandas. Zenodo. 10.5281/zenodo.3509134

Tonkin-Hill, G., Lees, J.A., Bentley, S.D., Frost, S.D., Corander, J. (2019). Fast hierarchical Bayesian analysis of population structure. Nucleic Acids Research, 47 (11), 5539–5549.

Tonkin-Hill, G., MacAlasdair, N., Ruis, C., Weimann, A., Horesh, G., Lees, J.A., … Parkhill, J. (2020). Producing polished prokaryotic pangenomes with the Panaroo pipeline. Genome Biology, 21 (1), 1–21.

van Kassel, M.N., Janssen, S.W., Kofman, S., Brouwer, M.C., van de Beek, D., Bijlsma, M.W. (2021). Prevalence of group b streptococcal colonization in the healthy non-pregnant population: a systematic review and meta-analysis. Clinical Microbiology and Infection, 27 (7), 968–980.

Viana, D., Comos, M., McAdam, P.R., Ward, M.J., Selva, L., Guinane, C.M., … Penadés, J.R. (2015). A single natural nucleotide mutation alters bacterial pathogen host tropism. Nature Genetics, 47 (4), 361–366.

