## Supplementary appendix for "Three Accessory Gene Clusters Drive Host-Adaptation in Group B *Streptococcus*"

### A Supplementary appendix: Methods

#### A.1 Whole genome sequencing

New genome sequence data was generated for n=91 isolates at two institutions. The University of Glasgow had n=44 isolates (originating from Vietnam) sequenced through MicrobesNG (Birmingham, UK). Isolates were plated on sheep blood agar (E&O Laboratories, Bonnybridge, UK) and grown overnight at 37°C to confirm viability and purity. One colony of each isolate was inoculated into Todd-Hewitt broth (Oxoid - Thermo Fisher Scientific, Waltham, Massachusetts, US) and incubated aerobically at 37°C overnight. DNA was extracted with the GenElute Bacterial Genomic DNA Kit (Sigma-Aldrich, St. Louis, Missouri, US) as per the manufacturer's instructions. Library preparation was carried out with the Nextera XT DNA Sample Preparation Kit and Miseq Reagent Kit V2 Library Preparation Kit (Illumina Inc., San Diego, California, US) and DNA was sequenced on an Illumina HiSeq platform. Similarly, Illumina sequencing was performed by the Genome Institute of Singapore (GIS) on n=47 isolates (originating from Vietnam and Thailand) with the Efficient Rapid Microbial Sequencing (GERMS) platform (https://www.a-star.edu.sg/gis/our-science/genome-architecture-and-design/germs).

#### A.2 Genome selection

The initial dataset comprised 2,437 GBS genomes, including newly generated data (n=91, see section A.1) and data from public repositories (NCBI and ENA, literature was reviewed in March 2020). When data were available in the form of raw reads (n=2,047 genomes), genomes were assembled with either SPAdes v3.14.0 (Bankevich et al., 2012) (n=1,025) or velvet v1.2.10 (Zerbino & Birney, 2008) (n=1,022), the latter used as standard in the Sanger Institute assembly pipeline.

We subsequently: i) curated metadata from the literature; ii) ran multi-locus sequence typing (MLST) with SRST2 v0.2.0 (Inouye et al., 2014) and run_MLST.py (the former on raw reads, the latter in case of assembled genomes)[1]; iii) determined the capsular type with blastn v2.12.0+ (Camacho et al., 2009) based on the method developed by Metcalf et al. (2017), and confirmed dubious cases with that of Sheppard et al. (2016); iv) de-duplicated the dataset for clonal isolates with the drop_duplicates() method within Pandas v1.1.3 (McKinney, 2011), based on the following data: country (down to the province/state level or city for countries such as Canada and the USA), host-species (including specific fish species e.g., tuna, tilapia etc.), origin/clinical manifestation (e.g., carriage vs invasive), year of isolation, farm/unit of origin (this was applicable to most bovine and fish isolates, but only to a few human isolates for which herds-persons had been sampled together with their livestock). A total of 1,158 genomes were eliminated at this step, including all genomes from Kawasaki et al. (2018) (n=39), because of inconsistencies between the isolate id/metadata and the associated genome assemblies for some genomes (e.g., capsular type and ST were not matching).

We then tested the remaining 1,279 genomes for assembly quality with QUAST v5.0.2 (Gurevich, Saveliev, Vyahhi, & Tesler, 2013). Reference ranges for GC content (%) and total genome length (bp) were calculated as their mean (35.38 and 2,074,536, respectively) $\pm$ twice the standard deviation (2SD) (reference ranges for GC content: 34.63-36.13; total length: 1,627,943-2,521,130). Genomes that were outside at least one of these ranges (n=25) were omitted from further analyses.

The resulting final dataset comprised 1,254 genomes representative of 9 host groups (bovine n=342; camel n=17; dog n=6; fish n=180, for some of which the species was available; food market fish samples n=26; frog n=4; goat n=2; human n=671; sea mammals n=6), across 7 continents or regions (South America n=431; Europe n=410; North America n=184; Asia n=163; Africa n=40; Oceania n=19; Central America n=5) and spanning 7 decades (1953-2021).

[1] Single locus variants (SLV) of existing sequence types (ST) were detected (n=14; n=2 ST103, n=2 ST314, n=5 ST552, n=3 ST61, n=1 ST1409, n=1 ST415); of note, for n=75 genomes from Brazil ST could not be determined because a *glcK* allele could not be assigned (n=15 genomes from bovine, n=60 from fish)

#### A.3 Core genome analyses

To reconstruct a core genome phylogeny, the snippy-multi script from the snippy suite v4.4.5 (https://github.com/tseemann/snippy) was used for alignment. To define the best topology of the tree, an outgroup was included in a first analysis, including reference genomes from the closely related species *Streptococcus pyogenes* (NCTC8224, NCTC12059, NCTC12069, NCTC13737, NCTC13744). The snippy-clean_full_aln script was then run on the core.full.aln file, on which gubbins v3.2.0 (Croucher et al., 2014) was run to remove recombination. IQ-TREE v.2.0.6 (Nguyen, Schmidt, Von Haeseler, & Minh, 2015) was then used to infer the phylogeny on the filtered_polymorphic_sites file from gubbins, with a GTR model with 1,000 bootstrap replicates. The tree was visualised in Microreact (Argimón et al., 2016).

A clustering algorithm, fastbaps v1.0.8 (Tonkin-Hill, Lees, Bentley, Frost, & Corander, 2019), was run on the same filtered_polymorphic_sites file with the optimise.symmetric parameter to define genomic clusters. These were then renamed, inheriting numbers from the 7-gene MLST nomenclature of the most represented ST within each population, preceded by the acronym SL for sub-lineage. For example, population 1 was mostly represented by ST17 isolates, and it was therefore named SL17 (Fig. C.2). CG were identified manually on the phylogeny within each SL.

For the classification of SL and CG in host specialists, host-adapted, and host generalists, the prevalence within SL/CG of the different hosts was calculated (Fig. 1B). The prevalence in the dominant host within each SL/CG was then plotted as a histogram, to identify cut-offs. Host specialists include those SL/CG that primarily occur in one host species or host group (e.g., SL552 for fish, SL61 and SL91 for bovine, SL22 for human, SL609 and SL612 for camel), although rare exceptions of their isolation from other hosts may exist in the literature (e.g., incidental report of SL61 in humans (Li et al., 2018)). Host-adapted SL/CG were defined as having more than 80% but less than 98% of isolates originating from the same host species or host group; and host generalists, were defined as having no more than 80% of isolates originating from a single host or host group (Fig. C.1).

Homologous recombination was visualised with Phandango (Hadfield et al., 2018) from the gubbins-generated recombination-prediction file. A generalised linear model was run in RStudio v2022.07.01, R v4.2.0 (2022-04-22), on the output from gubbins, after having mapped the internal nodes and the leaves to the corresponding CG, to test association between the host-specialisation level (generalists, adapted, specialists) and the ratio of recombination vs mutation (r/m).

The total number of CDS per genome was assessed with the panaroo-qc script.

#### A.4 Accessory genome analyses

To calculate pairwise distances of accessory gene content of isolates (using the Jaccard similarity index), a gene presence/absence matrix was generated with panaroo v1.2.0 from gff files annotated with Prokka v1.14.5 (Seemann, 2014). The matrix was processed with GraPPLE (Graphical Processing for Pangenome Linked Exploration) (downloaded on 27th April 2021; https://github.com/JDHarlingLee/GraPPLE). The resulting file was visualised with Graphia v2.2 (Freeman et al., 2022) (k-NN using edge weight k=18 and descending rank order).

Pilus island genes (Ap1 and Ap2, for both PI-1, PI-2a and PI-2b), which are important surface proteins, were detected with tblastn. This was done because an association between pilus variant and host of origin was reported in GBS (Lyhs et al., 2016), suggesting that pili may play a role in host adaptation, with PI-2a being the most common variant in human isolates and PI-2b the most frequent among bovine isolates.

#### A.5 Genome-Wide Association Studies (pyseer and scoary)

For the pangenome-based method, starting from the panaroo-generated presence/absence gene matrix, scoary v1.6.16 (Brynildsrud, Bohlin, Scheffer, & Eldholm, 2016) was used to detect gene-enrichment in the three host groups (human n=671, bovine n=342, fish n=180). This was done with the no_pairwise flag, as described in the scoary GitHub page (https://github.com/AdmiralenOla/Scoary). Sensitivity, specificity, *p*-values and negative/positive correlation of gene variants with host groups were assessed.

For the unitig-based method, the pyseer suite v1.3.3 (Lees, Galardini, Bentley, Weiser, & Corander, 2018) was used to find association between unitigs and host groups with a mixed effects model. Unitig-counter v1.0.5 was used to count unitigs on the 1,254 genome assemblies of the deduplicated dataset. The phylogeny_distance.py script was used to create a distance matrix from the core genome phylogeny. A binary (0 vs 1) phenotype file was created for each of the three host groups, and pyseer with a linear mixed model was run for each group (e.g., pyseer –lmm –phenotypes host.pheno –kmers unitigs_clean.txt.gz –similarity phylogeny_K.tsv –output-patterns host_unitigs_patterns.txt –cpu 24). The count_patterns.py script was then used to determine a significance threshold using the number of unique unitig patterns (patterns: 260,474; threshold: $1.92 \times 10^{-7}$). Significant unitigs were then extracted and mapped to reference genomes (NZ_CP019979 for human, CP008813 for bovine, CP003919 for fish) and visualised with Phandango (Hadfield et al., 2018). Manhattan plots were built with seaborn v11.2 (Waskom, 2021). Unitigs were then annotated with annotate_hits_pyseer with host-specific references (complete genome assemblies available from NCBI): NZ_CP019979 for the human phenotype (an ST19 serotype III isolate), CP008813 for the bovine phenotype (an ST103 serotype Ia isolate; one of the only two complete GBS genomes from bovine that are publicly available) and CP003919 for fish (an ST552 serotype Ib). The former two were plotted with python (Fig. C.3A and Fig. C.3B). Pyseer also allows unitig annotations from a list of reference and draft genomes, which was done for the three phenotypes (human: NZ_CP010867, NZ_CP013908, NZ_CP011329, NZ_CP019979, NZ_CP007570, NZ_CP007571, NZ_CP007572, NZ_CP020449, CP010319, CP020387, NZ_CP010874, NZ_CP010875, NZ_CP019978; bovine: CP008813; ERS1796640 [draft], JN_BR_Sag015 [draft], JN_BR_GBS104 [draft]; fish: CP003919, CP007482, CP019802, NC_018646, STIR-CD-25); these did not vary the results from annotation with one reference.

#### A.6 *In vivo* assessment of the role of Locus 3 in fish infection

##### A.6.1 Bacterial strains and plasmids

Strains and growth media are shown in Supplementary Tab. C.1. The temperature-sensitive *Escherichia coli* shuttle vector pG$^+$host9 (Maguin, Prevost, Ehrlich, & Gruss, 1996) was used for plasmid mediated allele exchange mutagenesis in GBS strains to generate ST7 and ST283 knock-out mutants of Locus 3 (ΔLocus3). Genomic DNA and plasmid DNA were extracted and purified using a GenElute bacterial genomic DNA extraction kit (Sigma-Aldrich) and a QIAprep Spin Miniprep kit (Qiagen), respectively. DNA was quantified using NanoDrop One UV-Vis spectrophotometer (ThermoFisher Scientific). All kits were used according to manufacturers' guidelines.

##### A.6.2 Construction of the pG$^+$host9 mutagenesis plasmid

Genomic sequences flanking the mutagenesis target regions were amplified from GBS ST7 gDNA using the Phusion PCR kit (New England Biolabs - NEB). Oligonucleotide primers for amplifying 5' and 3' flanking PCR sequences are described in Supplementary Tab. C.2. PCR products were visualised using 1% (w/v) TAE agarose gel electrophoresis, against a 1kb DNA ladder. Flanking PCR products were purified using a Qiaquick PCR purification kit (Qiagen). Purified flanking PCR products were digested with the appropriate *Bam*HI or *Eco*RI restriction endonuclease. Digested 5' and 3' flanking PCR products were ligated together using T4 DNA ligase to create mutant alleles. Mutant allele ligation products were amplified by Phusion PCR (NEB), using mutagenesis target primers F1 and R4, before Qiaquick PCR purification (Qiagen). The mutagenesis plasmid, pG$^+$host9, was digested with *Sma*I restriction endonuclease and purified with the Qiaquick PCR purification kit. Purified mutant alleles were ligated into linearized pG$^+$host9 using T4 DNA ligase with a molar insert to vector ratio of 10:1 and incubated overnight (16°C). Electrocompetent *E. coli* TG1-dev cells were prepared (Froehlich & Scott, 1991) and transformed with 4$\mu$L of overnight ligation mixture by electroporation using a 0.2mm electroporation cuvette and Gene Pulser XCell electroporation system (Bio-Rad) with parameters set at 2,000V; 25$\mu$F; 200$\Omega$. Transformed cells were immediately suspended in 400$\mu$L of SOC medium and incubated (37°C, 1 hour). Ten-fold dilutions of transformed cells were plated onto Luria-Bertani (LB) agar supplemented with 400$\mu$g/mL erythromycin and incubated (37°C, 2 days). *E. coli* TG1-dev transformants were screened for presence of mutant allele (mutagenesis primers F1-R4) by colony PCR using GoTaq Green mastermix. Plasmids were extracted from positive transformants and the mutant allele region was verified by sequencing using mutagenesis primers F1 and R4.

##### A.6.3 Transformation of GBS and temperature-mediated allele exchange mutagenesis

Electrocompetent GBS cells were prepared (Framson, Nittayajarn, Merry, Youngman, & Rubens, 1997) and transformed with 1$\mu$g of pG$^+$host9 construct by electroporation using a 0.1mm electroporation cuvette and Gene Pulser XCell electroporation system with parameters set at 2400V; 25$\mu$F; 100$\Omega$. Transformed cells were immediately suspended in 400$\mu$L of SOC medium and incubated (28°C, 1 hour). Transformed cells were plated neat onto TSA+2$\mu$g/mL erythromycin and incubated (28°C, 2 days). GBS transformants were screened for presence of mutant allele (mutagenesis primers F1-R4) by colony PCR using GoTaq Green mastermix. Plasmids were extracted from positive transformants, digested with *Bam*HI restriction endonuclease and visualised on a 1% TAE agarose gel against a 1kb DNA ladder for verification of mutagenesis plasmid constructs. Transformants containing mutagenesis plasmid constructs were cultured at 28°C (permissive temperature) overnight in 5mL of Tryptic Soy Broth (TSB) +2$\mu$g/mL erythromycin. Overnight cultures were diluted in 20mL TSB+2$\mu$g/mL erythromycin and grown at 28°C until reaching OD600$_{\text{nm}}$ 0.25, transferred to 37°C (non-permissive temperature) and incubated overnight. Ten-fold serial dilutions were prepared, 100$\mu$L of $10^{-1}$ to $10^{-4}$ dilutions was plated onto TSA+2$\mu$g/mL erythromycin and incubated (37°C, 2 days). Ten colonies of GBS were collected onto a single sterile inoculation loop and used to inoculate 20mL of TSB+2$\mu$g/mL erythromycin which was incubated (37°C, overnight). Overnight cultures were diluted 1:50 to TSB without antibiotic and incubated (28°C, until the end of the day), then diluted 1:50 to TSB without antibiotic and incubated again (28°C, overnight). This procedure was repeated once before ten-fold serial dilutions were prepared and 100$\mu$L of $10^{-4}$ to $10^{-7}$ dilutions plated onto TSA without antibiotic and incubated (37°C, overnight). 200 colonies per mutant strain were replica plated onto TSA+2$\mu$g/mL erythromycin and TSA without antibiotic and incubated (37°C, overnight). Replica GBS strains which were susceptible to erythromycin were selected from the TSA without antibiotic plate and incubated in TSB (37°C, overnight). Genomic DNA was extracted from overnight cultures. Mutagenesis of target Locus 3 regions were confirmed firstly by PCR amplification of gDNA using mutagenesis primers F1 and R4 and secondly by DNA sequencing across the gDNA target region with mutagenesis primers F1 and R4. Specific Locus 3 genes (galactokinase; $\alpha$-galactosidase; ABC sugar permease substrate binding protein; PTS system galactitol specific IIC component gene number 14) were screened using PCR to detect presence or absence in mutant strains using primers shown in Supplementary Tab. C.2.

##### A.6.4 Tilapia challenge studies

Challenge studies were conducted by a contract research organisation specialised in aquatic animal health (Ictyodev, France). Studies were performed to Good Laboratory Practice standards, approved by an internal Ethics Committee and the French Ministry of Research, and audited by an independent Quality Assurance partner. Challenge studies were conducted with wild type (WT) GBS ARG/BAC/2014-107 (ST7) and ARG/BAC/2016-6 (ST283) and their ΔLocus3 knock-out mutants.

Nile tilapia (*Oreochromis niloticus*) fingerlings were acclimatised for 1 week pre-challenge in 450L freshwater tanks at 30°C. In preparation for challenge, fish were transferred into 100L experimental tanks (n=4 replicates per GBS strain, n=2 negative control replicates) under the same ecological conditions. Ten fish from each experimental tank were challenged by intraperitoneal (IP) injection with 0.1mL of PBS (negative control) or 0.1mL of $10^5$ CFU/mL of GBS in PBS. IP-challenged tilapia were co-habited with 40 healthy tilapia per tank to provide contact challenge. Mortality of IP-challenged and contact-challenged fish was monitored for 21 days. Moribund fish were euthanized and included in mortality counts. Kidney and brain samples were taken from up to 10 fish per tank (n=5 IP-challenged, n=5 contact exposed) and used to check for presence of GBS, cultures of which were stored in BHI with 30% glycerol (v/v) at -80°C. Locus 3 gene specific PCR assays were performed to confirm the identity of strains isolated from challenged fish.

##### A.6.5 Statistical analysis

Kaplan-Meier survival analysis (May, 2009) was conducted to compare survival of knock-out mutants and their isogenic WT strains. A statistical Log-rank test was conducted (Kleinbaum & Klein, 2012), using Microsoft Excel software, to compare survival curves of tilapia challenged by WT vs knock-out mutants.

#### A.7 Time-scaled phylogenies of CG283 and SL23 and number of host-jumps

BEAST v2.6.6 (Bouckaert et al., 2014) was used to estimate the time of emergence of two clones of public health interest, CG283 and SL23. Only genomes with available isolation dates were included in the analyses (n=102 and n=187, respectively). Recombination-free alignment files were generated with snippy and gubbins. The best model parameters were chosen after carrying out tests on combinations of: i) substitution model HKY or GTR+G4, ii) strict clock and relaxed log-normal clock; iii) coalescent constant and coalescent Bayesian Skyline population. XmL files generated with BEAUTi were run in triplicate with 200 million generations, sampling frequency of 10,000 (burn-in 10%) and maximum likelihood estimation (MLE) with path stepping stone for model test. The best model for both was: GTR+G4 with a strict clock and a constant coalescent population size. Resulting trees from the tree replicate runs were combined with LogCombiner, and the final tree files were obtained with TreeAnnotator and visualised with FigTree v1.4.4 (http://tree.bio.ed.ac.uk/software/figtree/) and Microreact (Argimón et al., 2016).

### B Supplementary appendix: Results

#### B.1 Core genome analyses

Different ratios of recombination vs mutation (r/m) were observed among CG, with CG609 and CG17 showing the lowest values (n=0.0962 and n=0.139, respectively), and CG103, CG19 and CG817 the highest (n=5.87, n=5.83, and n=4.57, respectively) (Fig. 1D). However, no significant difference of r/m between the three groups of host-specialism level was observed (t-value generalists=11.632, adapted=-0.935, specialists=-1.123, p-value=0.4393).

#### B.2 Accessory genome analyses

Similar to previous studies (Lyhs et al., 2016), in our dataset we found a higher prevalence of PI-2a in human genomes (79.3%) compared to bovine (46.5%) and fish (30.6%) genomes. However, we found only a slightly higher frequency of PI-2b in bovine (53.2%) compared to human (20.9%), with the highest prevalence among fish genomes (70.6%). When observing the distribution of pili among the phylogeny (section B.4), it is clear that the association of these mobile elements is more associated with SL/CG than with the host of origin.

#### B.3 Genome-wide association studies (GWAS) - additional results

##### B.3.1 Scoary

For the human phenotype, together with positively-associated genes with high sensitivity and specificity (part of the *scpB* transposon, SE 91.1-92.6%, SP 57.3-72.2%), some high-scoring genes were negatively-associated, in particular genes belonging to the Lac.2 operon *lacEFG* (SE 5.3%, SP 50.9%). Ten genes were identified as human-specific by scoary, i.e., found uniquely in GBS genomes from humans, although not in all of them. This included a TetR transcriptional regulator and a multidrug transporter (see Supplementary file S2).
For the bovine phenotype, a toxin-antitoxin system (*pezAT*) was positively-associated (*pezA*: *p*-value $2.12 \times 10^{-92}$, SE 68.4%, SP 90.7%; *pezT*: *p*-value $1.75 \times 10^{-71}$, SE 59.9%, SP 90.8%). In addition, phage-related genes and genes belonging to an integrative-conjugative element (ICE), such as an LPxTG motif, also scored highly. These genes were positively-associated with the bovine phenotype (see section B.3.2). Bovine-specific genes (n=118) included transporters for antimicrobials (macrolides and fluoroquinolones) and bacteriocins (lactococcin, bacitracin), transposases (IS*Sag*8, IS4) and phage-associated genes (see Supplementary file S3).
Seventy-one genes were identified uniquely in the fish phenotype, including

bacteriocin precursors and transporters (bacitracin transporter, lactacin subunit) and transposases (IS*Rgn*1, IS*Stin*5, IS*Sag*8, IS*Stso*1). Twelve of these genes belonged to fish-specific loci identified by Delannoy et al. (2016) (locus 1, 2, 4, 5 6 and 8) (see Supplementary file S4).

##### B.3.2 Pyseer

The toxin-antitoxin system *pezAT* identified by scoary as significantly-associated with the bovine phenotype was also detected by pyseer, as were several genes belonging to an ICE (see section B.3.1) and a prophage. These same phage genes matched those of two incomplete prophages (which show a high similarity to each other) that had previously been reported as bovine-associated by Richards et al. (2011) (the prophage in reference genome CP008813 had 96.36% ID and 68% query coverage with phage A from Richards et al. (2011), and 93.70% ID and 67% query coverage with phage B) (see section B.3.3).
For the fish phenotype, pyseer results were unsatisfactory. Although population structure should have been controlled for (this was done after several iterative trials, setting an optimal minor allele frequency, or MAF, of a minimum of 0.04 and a maximum of 0.96), no evident peaks were present in Manhattan plots of any complete genome sequences from fish in this dataset (n=39).

##### B.3.3 BLAST

To confirm association of *pezAT*, the ICE and the prophage with the bovine phenotype, tblastn searches (thresholds of 90% identity and 80% query coverage) were performed on amino-acid sequences of *pezAT*, of two ICE genes (LPXTG cell wall anchor protein and N-6-DNA methylase) and two prophage genes (phage tail spike protein and phage tail tape measure protein). All three elements were confirmed as more prevalent among bovine GBS genomes, although not as common as Lac.2, and with variable prevalence in the other two host groups. *pezAT* in bovines: 46.5%; in humans: 5.4%; and in fish: 0%. ICE in bovines: 52.6% and 52.3% (LPXTG and N-6-DNA-methylase, respectively); in humans: 27.0% and 27.1%; and in fish: 10.6% for both (in fish, ICE genes were uniquely found in ST7 genomes). Prophage in bovines: 69.0% and 69.9% (phage tail spike protein and phage tail tape measure protein, respectively); in humans: 37.9% and 38.5%; and in fish: 10.6% for both (in fish, prophage genes were uniquely found in ST7 genomes, the same ones that also carried ICE genes).

#### B.4 Group B *Streptococcus* sublineages

### B.4.1 SL17

Sublineage 17 (SL17) (Fig. C.4A), is a human-adapted lineage that shows rare possibility of spill-over events in both cattle and fish. The most prevalent capsular type (CPS) detected in this lineage is III, with a few CPS IV being detected. Strong human-association of this lineage is reflected in the pattern of presence/absence of the three host-associated accessory gene clusters: *scpB*+, Lac.2- (with two exceptions) and Locus 3-. Pilus island genes associated with this lineage are PI-1 and PI-2b. Recombination is almost absent from this SL (Fig. 1A).

### B.4.2 SL19

Sublineage 19 (SL19) (Fig. C.4B), is a human-adapted lineage that shows possibility of spill-over events, primarily in cattle. Two main capsular types are detected in this lineage, whereby CPS II is associated with CG28, and CPS III with CG19. CG19 includes six serotypes, with relatively high prevalence of CPS V in separate branches, indicative of multiple independent capsular switching events. Human-association of this lineage is reflected in the pattern of presence/absence of the three host-associated accessory gene clusters: *scpB*+, Lac.2- (with a few exceptions) and Locus 3-. Pilus island genes associated with this lineage are PI-1 and PI-2a. Recombination is present in SL19, some of which is shared with SL183 and SL1, but lineage-specific recombination blocks can also be observed (Fig. 1A).

##### B.4.3 SL91 and SL61

Sublineages 91 and 61 (SL91 and SL61) (Fig. C.5A), are bovine-specialists. One main CPS, II, is associated with SL61, whereas SL91 is associated almost equally with CPS II and III. Capsular switching is observed in both SL, with one branch showing acquisition of CPS Ia. All isolates on this branch originate from Colombia. Bovine-specificity of these lineages is reflected in the pattern of presence/absence of the three host-associated accessory gene clusters: *scpB*- (with one exception), Lac.2+ and Locus 3-. Pilus island genes associated with these lineages are primarily PI-2b, with four genomes of SL91 also carrying PI-1, a combination otherwise mostly seen in SL17. All SL91 isolates with this PI profile originate from Italy. Recombination is present in SL61 and SL91, but it does not involve large recombination events, and it is mostly lineage-specific (Fig. 1A).

##### B.4.4 SL103 and SL314

Sublineages 103 and 314 (SL103 and SL314) (Fig. C.5B), are host-generalist lineages with a tendency towards bovine-association. Most genomes carry CPS Ia, except for a sub-clade of CPS III, which is mostly associated with human isolates. The lack of a strong host association is reflected in the mixed pattern of presence/absence of Lac.2. Surprisingly, *scpB* is mostly absent, despite isolation of SL103 and SL314 from humans, whereas Locus 3 is commonly present, even though this lineage is not represented among available fish GBS genomes. Pilus island genes associated with these lineages are primarily PI-2b. Recombination is almost absent from these two SL (Fig. 1A).

### B.4.5 SL552

Sublineage 552 (SL552) (Fig. C.6A), is restricted to fishes and frogs. Although sometimes described as a poikilothermic lineage Richards et al. (2019), it has not been reported from reptiles whereas other types of GBS have. Genomes in SL552 all carry CPS Ib (in one case the *in silico* CPS screening gave two best possible matches: Ib and III), and fish-specificity of this lineage is reflected in the pattern of presence/absence of the three host-associated accessory gene clusters: *scpB*-, Lac.2- and Locus 3+. Pilus island genes associated with this lineage belong to PI-2b, although PI-2bAP2 is absent from CG261. Recombination is absent from the entire sublineage (Fig. 1A).

### B.4.6 SL1

Sublineage 1 (SL1) (Fig. C.6B), is a host-generalist lineage. Most genomes of CG1 carry CPS V, those of CG459 have serotype IV, whereas several CPS are associated with CG817 (including Ia, II, III, IV and VI). The latter CG is shows the greatest diversity within SL1, with long branches, variable association with humans or cattle, variable pattern of presence/absence of host-associated genes, of pili (Fig. C.6B), and lineage-specific recombination blocks (Fig. 1A).CG459 shows the lowest diversity, as reflected in branch length, host-association (primarily human) and presence/absence of host-associated genes (mostly *scpB*+, Lac.2- and Locus 3-). CG1 is associated with both cattle and humans, and interspecies transmission is likely to be possible, especially for those isolates that carry both *scpB* and Lac.2. Pilus island genes associated with CG1 belong predominantly to PI-1 and PI-2a. Recombination is widely present in SL1, which shares some recombination blocks with SL19 and SL283, whereas other blocks are lineage-specific 1A).

### C Supplementary appendix: Fig.s and Tab.s

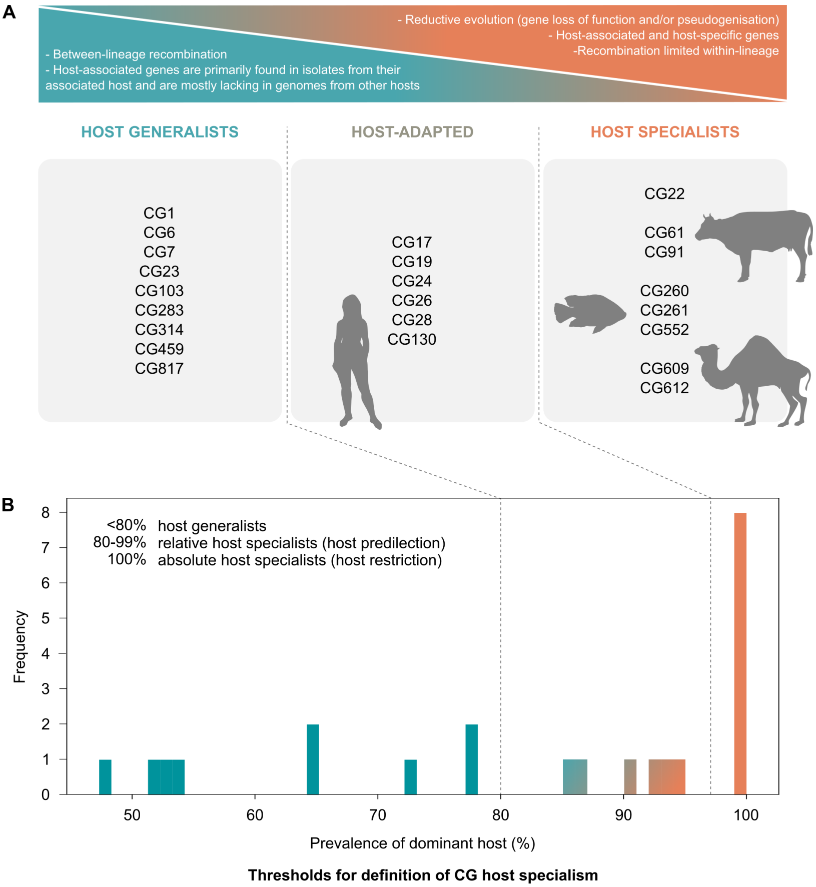


**Fig. C.1** Diagram illustrating host specialism levels in Group B *Streptococcus* (GBS) clonal groups (CG). A) Host generalist lineages show extensive between-lineage homologous recombination and the host associated accessory genes (*scpB*, Lac.2, Locus 3) are primarily found in isolates from their associated host, while they are lacking from isolates from other hosts (e.g., Lac.2 tends to be present in most isolates from cattle and absent from most isolates from humans). Host specialist lineages (with or without host restriction) can be associated with reductive evolution (e.g., gene loss of function and genome reduction as in SL552, which comprises CG260, CG261 and CG552), they carry host-associated genes (e.g., CG61 and CG91 all carry Lac.2, SL552 all carry Locus 3) and they either show absence of recombination (SL552) or recombination limited within CG. Host-adapted CG are primarily associated with one host and they show all characteristics of the host-specialist lineages except for genome reduction and pseudogenisation. B) Thresholds used for the categorisation of CG in the three levels of host specialism; the prevalence in the dominant host for each CG was plotted to identify cut-offs.

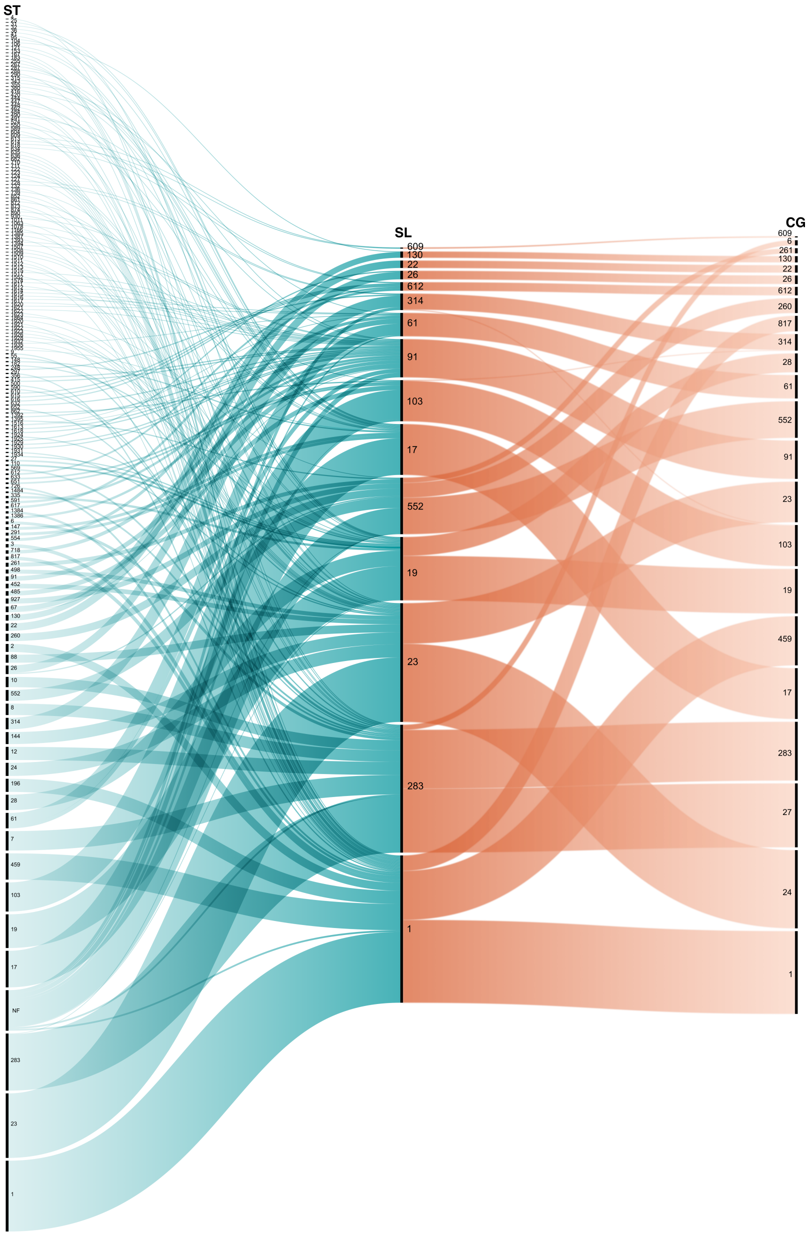


**Fig. C.2** Alluvial diagram showing the inheritance of 7-gene multi-locus sequence typing (MLST) nomenclature on sublineages (SL) and clonal groups (CG). Note that some new sequence types (ST) and those that could not be defined due to a missing locus are indicated with NF (not found).

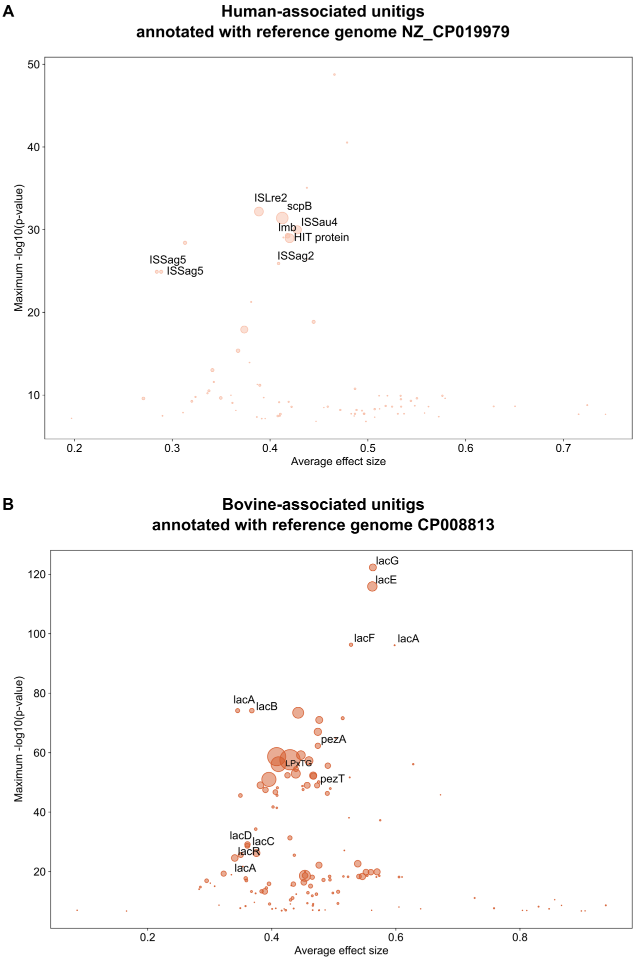


**Fig. C.3** Annotated unitigs mapped to two reference genomes. A) Human-associated unitigs annotated to reference genome NZ_CP019979; B) Bovine-associated unitigs annotated to reference genome CP008813. Genes in common with results from scoary have been labelled. For the human-associated unitigs, labelled genes are all part of the *scpB* transposon. For the bovine-associated unitigs, *lacABCDEFGR* are all part of the Lac.2 operon, whereas *pezAT* are associated with an integrative conjugative element.

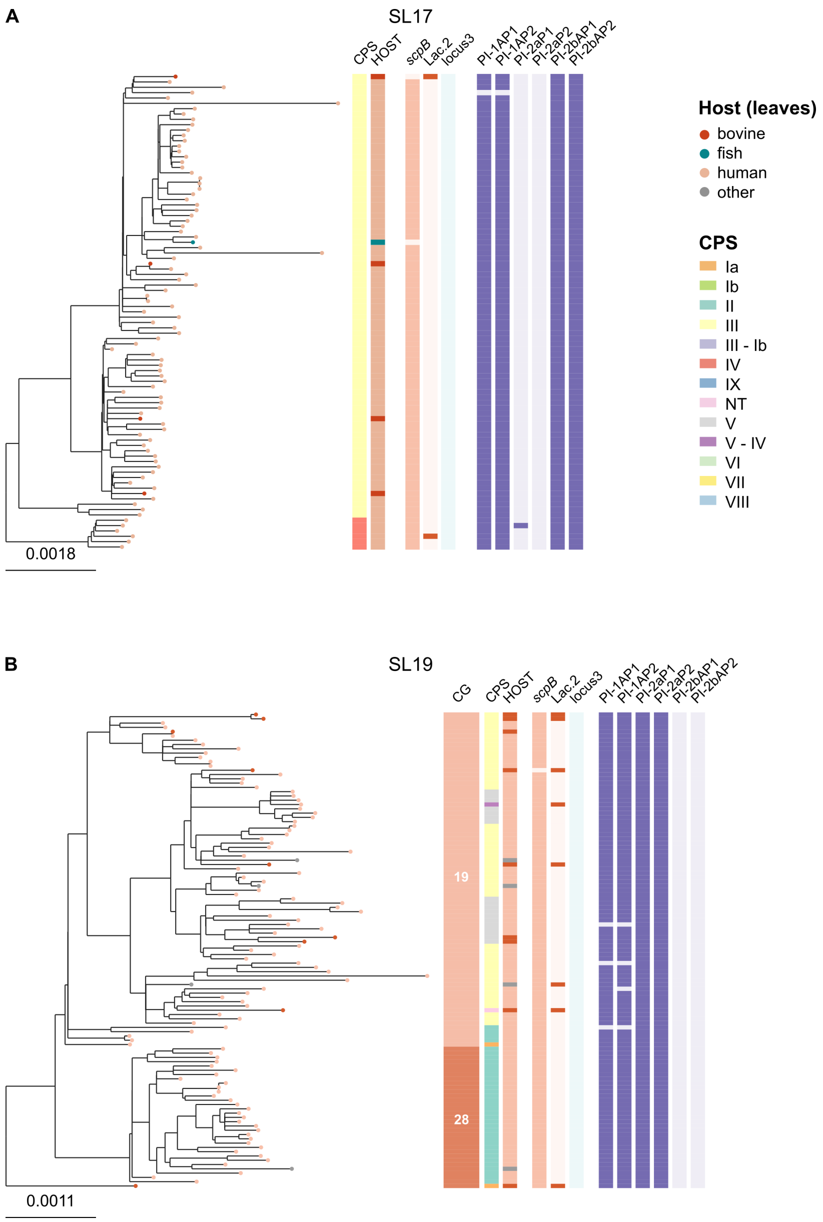


**Fig. C.4** Maximum-likelihood phylogenetic tree of SL17 (A), and of SL19 (B). Capsular type (CPS) and host of origin are indicated. Presence/absence of host-associated accessory genes (*scpB*, Lac.2, Locus 3) and pilus island genes is shown. In SL19, clonal groups (CG) are also indicated; this was not done for SL17, as it comprises one unique CG (CG17). SL17 and SL19 are human-specialists, and they show a complete lack of Locus 3 and a preponderant absence of Lac.2, whereas the human-associated gene *scpB* is highly conserved in both SL. Pilus islands PI-1 and PI-2b are almost inariably present in SL17, whilst this combination is rare in other sublineages.

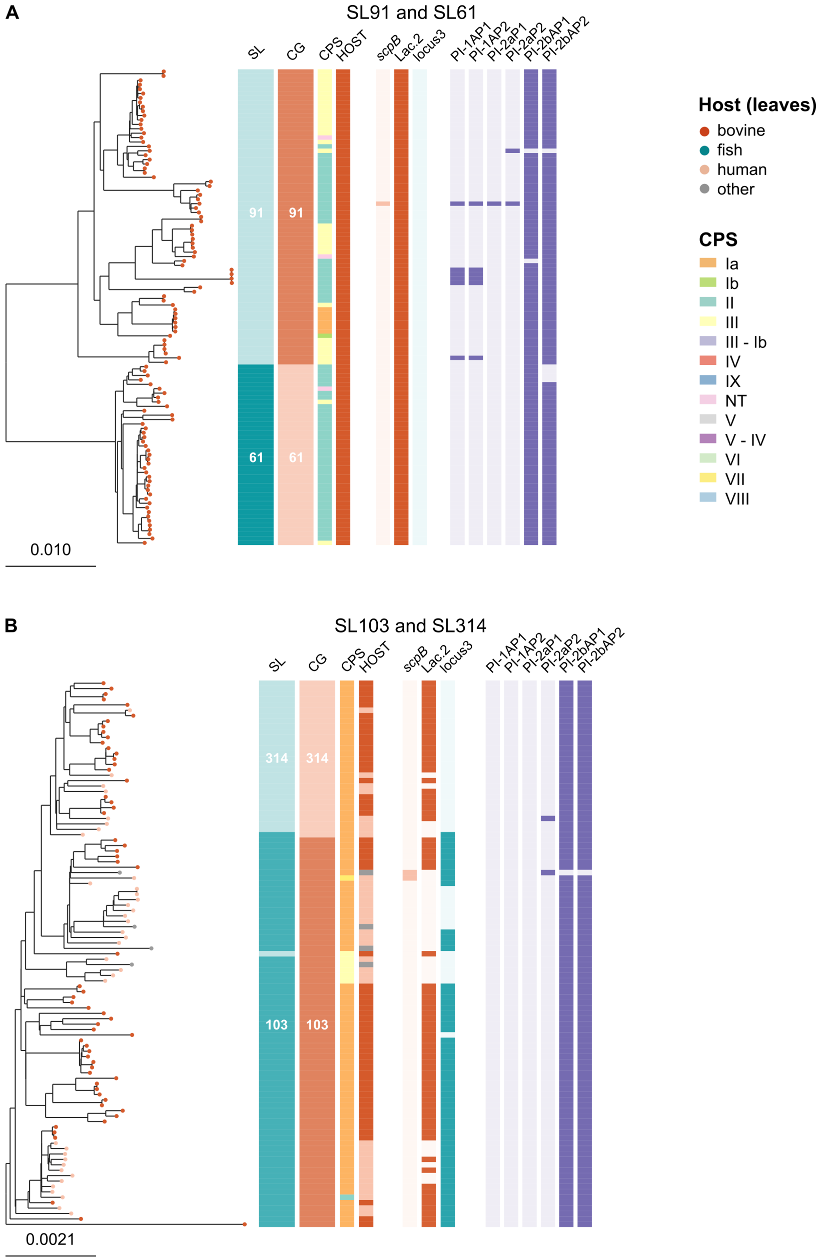


**Fig. C.5** Maximum-likelihood phylogenetic tree of SL91 and SL61 (A), and of SL103 and SL314 (B). Sublineage (SL), clonal group (CG), capsular type (CPS) and host of origin are indicated. Presence/absence of host-associated accessory genes (*scpB*, Lac.2, Locus 3) and pilus island genes is shown. SL91 and SL61 are bovine-specialists. They show a total absence of Locus 3 and an almost complete absence of *scpB*, whereas the bovine-associated Lac.2 gene cluster is highly conserved across both SL. Pilus island PI-2b is found in both lineages and PI-1 has been acquired in one geographically restricted (Italy) branch of SL91. Clustering of host-generalist lineages SL103 and SL314 was not as neat as for the other lineages (one genome phylogenetically close to SL103 was categorised as belonging to SL314 and vice-versa). Despite being commonly found in cattle and humans, *scpB* is largely absent whereas Lac.2 and Locus 3 are often present. Pilus island PI-2b is highly conserved across both SL.

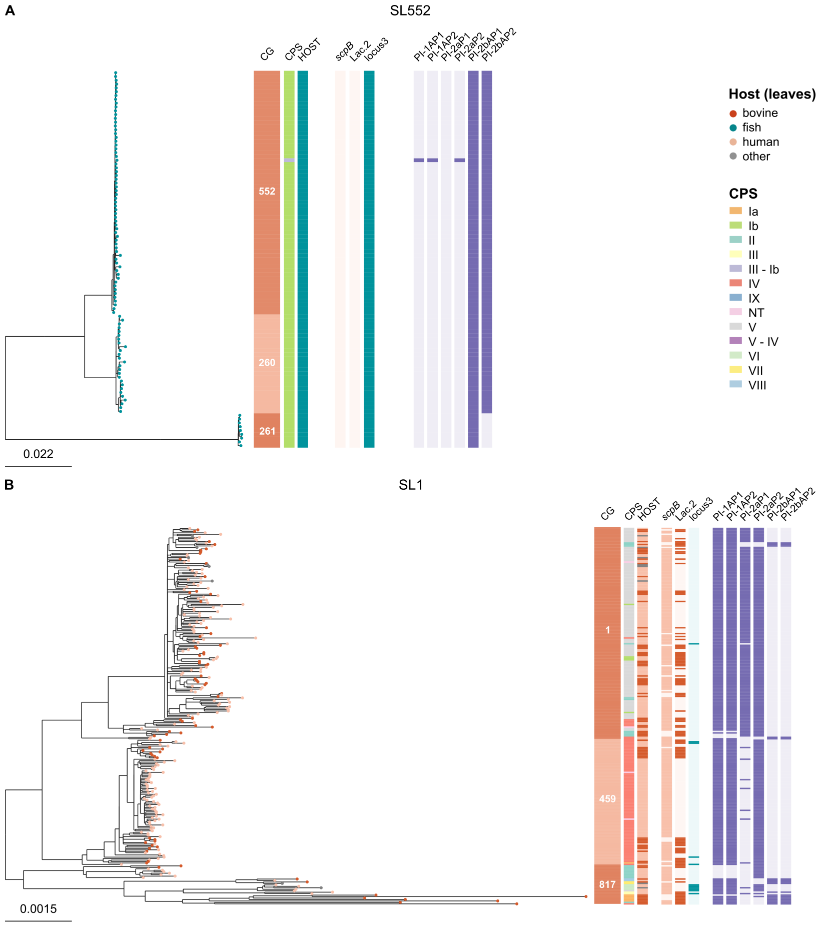


**Fig. C.6** Maximum-likelihood phylogenetic tree of SL552 (A), and of SL1 (B). Clonal group (CG), capsular type (CPS) and host of origin are indicated. Presence/absence of host-associated accessory genes (*scpB*, Lac.2, Locus 3) and pilus island genes is shown. SL552 is a fish-specialist (fish and other cold-blooded species such as frogs). It shows a complete absence of *scpB* and Lac.2, whereas Locus 3 is omnipresent. Pilus island PI-2b is found in all genomes from this SL, albeit incomplete in CG261. SL1 is a host-generalist (human, bovine, other hosts). *scpB* is highly prevalent in this lineage, whereas Lac.2 is variably present, and mostly associated with bovine isolates. Locus 3 is mostly absent. A tendency towards human-specialisation can be observed within CG459 (CPS IV), in which a sub-clade is uniquely associated with the human host, all genomes carry *scpB* and lack both Lac.2 and Locus 3. Predominant pilus island genes associated with CG1 are PI-1 and PI-2a, whereas CG459 and CG817 mostly show presence of PI-1 and elements of PI-2a. Across SL1, especially in CG817, some isolates carry PI-2b genes.

**Table C.1** Bacterial strains and culture conditions used to generate Group B *Streptococcus* mutants.

| Strain | Description | Culture medium | Culture temperature ($^{\circ}C$) | Antibiotic selection |
|---|---|---|---|---|
| ***E. coli*** | | | | |
| TG1-dev | Cloning host permissive for $pG^{+}$host9 plasmid replication at 37$^{\circ}$C | Luria-Bertani (LB) | 37 | none |
| TG1-dev $pG^{+}$host9 | Maintains empty $pG^{+}$host9 plasmid | Luria-Bertani (LB) | 37 | Erm 400$\mu$g/mL |
| TG1-dev $pG^{+}$host9:Locus3 | Cloning host maintaining Locus 3 mutagenesis plasmid construct | Luria-Bertani (LB) | 37 | Erm 400$\mu$g/mL |
| **Group B** ***Streptococcus*** | | | | |
| ARG/BAC/2014-107 | ST7 clinical isolate from a tilapia streptococcosis outbreak in Canada | Tryptic Soy Broth (TSB) | 37 | none |
| ST7 $pG^{+}$host9:Locus3 | ARG/BAC/2014-107 isolate maintaining entire Locus 3 mutagenesis plasmid construct | Tryptic Soy Broth (TSB) | 28 | Erm 2$\mu$g/mL |
| ST7 ΔLocus3 | ARG/BAC/2014-107 Locus 3 knock out mutant | Tryptic Soy Broth (TSB) | 37 | none |
| ARG/BAC/2016-6 | ST283 clinical isolate from a tilapia streptococcosis outbreak in South East Asia | Tryptic Soy Broth (TSB) | 37 | none |
| ST283 $pG^{+}$host9:Locus3 | ARG/BAC/2016-6 isolate maintaining entire Locus 3 mutagenesis plasmid construct | Tryptic Soy Broth (TSB) | 28 | Erm 2$\mu$g/mL |

**Table C.2** Oligonucleotide primers used for mutagenesis of Group B *Streptococcus*.

| Primer | Sequence (5'-3') | Annealing temperature ($^{\circ}C$) | Amplicon (bp) | Use |
|---|---|---|---|---|
| L3 F1 | TTCGCAAACGCTTCAAGGCTG | 71.4 | 471 | Locus 3 mutagenesis |
| L3 02_*Bam*HI | CGCGCGGGATCCTGAGCGAGCAGCAGATGCG | 71.7 | | |
| L3 03_*Bam*HI | CGCGCGGGATCCGCTGTCTAGCGTAGCGCC | 64 | 480 | |
| L3 R4 | AGGGCAGTTTGAGTGCATTTCC | 68 | | |
| galK Fwd | AAATCGGCAAGCAGACAGAAAATGAAT | 70 | 574 | Detection of Locus 3 galactokinase gene |
| galK Rev | GCAATAGCACAACCGCCAAAACC | 71.8 | | |
| $\alpha$-gal Fwd | AAGGGTGCTAGTAGTGCCGAACATAAT | 68 | 449 | Detection of Locus 3 $\alpha$-galactosidase gene |
| $\alpha$-gal Rev | AACCAGCCATCATCCATAACAAAAAGT | 68.6 | | |
| ABCsbp Fwd | ATGAAAAAAATGGGTTACTATATGGG | 62.6 | 1278 | Detection of Locus 3 ABC sugar permease substrate binding protein gene |
| ABCsbp Rev | TTATTTTTCCCGCTCAAGC | 61.7 | | |
| PTSIIC-14 Fwd | ATGGATGTTCTACAAACTGTTATTAAT | 58.9 | 1449 | Detection of Locus 3 galactitol specific PTS transporter IIC subunit (gene 14) |
| PTSIIC-14 Rev | TTATTCCACAATCAAATTCGT | 58.4 | | |

### References


Argimón, S., Abudahab, K., Goater, R.J., Fedosejev, A., Bhai, J., Glasner, C., ... others (2016). Microreact: visualizing and sharing data for genomic epidemiology and phylogeography. *Microbial Genomics*, *2*(11), e000093.

Bankevich, A., Nurk, S., Antipov, D., Gurevich, A.A., Dvorkin, M., Kulikov, A.S., ... Pevznernal, P.A. (2012). SPAdes: a new genome assembly algorithm and its applications to single-cell sequencing. *Journal of Computational Biology*, *19*(5), 455–477.

Bouckaert, R., Heled, J., Kühnert, D., Vaughan, T., Wu, C.-H., Xie, D., ... Drummond, A.J. (2014). Beast 2: a software platform for bayesian evolutionary analysis. *PLoS Computational Biology*, *10*(4), e1003537.

Brynildsrud, O., Bohlin, J., Scheffer, L., Eldholm, V. (2016). Rapid scoring of genes in microbial pan-genome-wide association studies with Scoary. *Genome Biology*, *17*(1), 1–9.

Camacho, C., Coulouris, G., Avagyan, V., Ma, N., Papadopoulos, J., Bealer, K., Madden, T.L. (2009). BLAST+: architecture and applications. *BMC Bioinformatics*, *10*(1), 421.

Croucher, N.J., Page, A.J., Connor, T.R., Delaney, A.J., Keane, J.A., Bentley, S.D., ... Harris, S.R. (2014). Rapid phylogenetic analysis of large samples of recombinant bacterial whole genome sequences using gubbins. *Nucleic acids research*, *43*(3), e15–e15.

Delannoy, C.M., Zadoks, R.N., Crumlish, M., Rodgers, D., Lainson, F.A., Ferguson, H., ... Fontaine, M.C. (2016). Genomic comparison of virulent and non-virulent *Streptococcus agalactiae* in fish. *Journal of Fish Diseases*, *39*(1), 13–29.

Framson, P.E., Nittayajarn, A., Merry, J., Youngman, P., Rubens, C.E. (1997). New genetic techniques for group b streptococci: high-efficiency transformation, maintenance of temperature-sensitive pwv01 plasmids, and mutagenesis with tn917. *Applied and Environmental Microbiology*, *63*(9), 3539–3547.

Freeman, T.C., Horsewell, S., Patir, A., Harling-Lee, J., Regan, T., Shih, B.B., … Angus, T. (2022). Graphia: A platform for the graph-based visualisation and analysis of high dimensional data. *PLoS Computational Biology*, *18*(7), e1010310.

Froehlich, B.J., & Scott, J.R. (1991). A single-copy promoter-cloning vector for use in *Escherichia coli*. *Gene*, *108*(1), 99–101.

Gurevich, A., Saveliev, V., Vyahhi, N., Tesler, G. (2013). QUAST: quality assessment tool for genome assemblies. *Bioinformatics*, *29*(8), 1072–1075.

Hadfield, J., Croucher, N.J., Goater, R.J., Abudahab, K., Aanensen, D.M., Harris, S.R. (2018). Phandango: an interactive viewer for bacterial population genomics. *Bioinformatics*, *34*(2), 292–293.

Inouye, M., Dashnow, H., Raven, L.-A., Schultz, M.B., Pope, B.J., Tomita, T., … Holt, K.E. (2014). SRST2: rapid genomic surveillance for public health and hospital microbiology labs. *Genome Medicine*, *6*(11), 90.

Kawasaki, M., Delamare-Deboutteville, J., Bowater, R.O., Walker, M.J., Beatson, S., Zakour, N.L.B., Barnes, A.C. (2018). Microevolution of *Streptococcus agalactiae* ST-261 from Australia indicates dissemination via imported tilapia and ongoing adaptation to marine hosts or environment. *Applied and Environmental Microbiology*, *84*(16), e00859–18.

Kleinbaum, D.G., & Klein, M. (2012). *Survival Analysis: A Self-Learning Text.* Springer, New York.

Lees, J.A., Galardini, M., Bentley, S.D., Weiser, J.N., Corander, J. (2018). Pyseer: a comprehensive tool for microbial pangenome-wide association studies. *Bioinformatics*, *34*(24), 4310–4312.

Li, L., Wang, R., Huang, Y., Huang, T., Luo, F., Huang, W., … Gan, X. (2018). High incidence of pathogenic *Streptococcus agalactiae* ST485 strain in pregnant/puerperal women and isolation of hyper-virulent human CC67 strain. *Frontiers in Microbiology*, *9*, 50.

Lyhs, U., Kulkas, L., Katholm, J., Waller, K.P., Saha, K., Tomusk, R.J., Zadoks, R.N. (2016). *Streptococcus agalactiae* serotype IV in humans and cattle, northern Europe. *Emerging Infectious Diseases*, *22*(12), 2097.

Maguin, E., Prevost, H., Ehrlich, S.D., Gruss, A. (1996). Efficient insertional mutagenesis in lactococci and other gram-positive bacteria. *Journal of Bacteriology*, *178*(3), 931–935.

May, W.L. (2009). Kaplan-Meier Survival Analysis. *Encyclopedia of Cancer* (p. 1590–1593). Springer, New York.

McKinney, W. (2011). pandas: a foundational Python library for data analysis and statistics. *Python for High Performance and Scientific Computing*, *14*(9), 1–9.

Metcalf, B., Chochua, S., Gertz Jr, R., Hawkins, P., Ricaldi, J., Li, Z., ... Beall, B. (2017). Short-read whole genome sequencing for determination of antimicrobial resistance mechanisms and capsular serotypes of current invasive *Streptococcus agalactiae* recovered in the USA. *Clinical Microbiology and Infection*, *23*(8), 574–e7.

Nguyen, L.T., Schmidt, H.A., Von Haeseler, A., Minh, B.Q. (2015). IQ-TREE: a fast and effective stochastic algorithm for estimating maximum-likelihood phylogenies. *Molecular Biology and Evolution*, *32*(1), 268–274.

Richards, V.P., Lang, P., Bitar, P.D.P., Lefébure, T., Schukken, Y.H., Zadoks, R.N., Stanhope, M.J. (2011). Comparative genomics and the role of lateral gene transfer in the evolution of bovine adapted *Streptococcus agalactiae*. *Infection, Genetics and Evolution*, *11*(6), 1263–1275.

Richards, V.P., Velsko, I.M., Alam, T., Zadoks, R.N., Manning, S.D., Pavinski Bitar, P.D., ... Stanhope, M.J. (2019). Population gene introgression and high genome plasticity for the zoonotic pathogen *Streptococcus agalactiae*. *Molecular Biology and Evolution*, *36*(11), 2572–2590.

Seemann, T. (2014). Prokka: rapid prokaryotic genome annotation. *Bioinformatics*, *30*(14), 2068–2069.

Sheppard, A.E., Vaughan, A., Jones, N., Turner, P., Turner, C., Efstratiou, A., … Seale, A.C. (2016). Capsular typing method for *Streptococcus agalactiae* using whole-genome sequence data. *Journal of Clinical Microbiology*, *54*(5), 1388–1390.

Tonkin-Hill, G., Lees, J.A., Bentley, S.D., Frost, S.D., Corander, J. (2019). Fast hierarchical Bayesian analysis of population structure. *Nucleic Acids Research*, *47*(11), 5539–5549.

Waskom, M.L. (2021). seaborn: statistical data visualization. *Journal of Open Source Software*, *6*(60), 3021. Retrieved from https://doi.org/10.21105/joss.03021

10.21105/joss.03021

Zerbino, D.R., & Birney, E. (2008). Velvet: algorithms for *de novo* short read assembly using de Bruijn graphs. *Genome Research*, *18*(5), 821–829.
